# Thioredoxin interacting protein (TXNIP), a redox regulator, mediates the EPAC-RAP1 signaling dependency of primary melanoma

**DOI:** 10.64898/2025.12.29.696903

**Authors:** Sireesh Kumar Teertam, Mithalesh K. Singh, Sarah Altameemi, Sonia Gude, Sushmita Roy, Ryan Rossman, Michael A. Newton, Daniel D. Bennett, Nihal Ahmad, Xiaodong Cheng, Vijayasaradhi Setaluri

## Abstract

Progression of cutaneous primary melanoma that arises from melanocytes leads to lethal metastatic disease. Molecular mechanisms that control the growth of primary melanoma in the skin and promote progression are not fully understood. Previously we showed that RAP guanine exchange factors EPAC1/2 (Exchange Protein Activated by cyclic AMP) promote the growth of primary melanoma and loss of dependency on EPACs is associated with metastatic progression. In this study, we show that EPACs are activated during malignant transformation of melanocytes, and chemical inhibition or genetic deletion of EPAC inhibits melanomagenesis in *Braf/Pten* mice. Low expression of EPAC mRNA and its effector RAP1-GTP protein in primary melanoma correlate with better recurrence-free survival. RNAseq analysis of matched primary and metastatic melanoma cells treated with an EPAC inhibitor showed that TXNIP, an important regulator of redox homeostasis, is a downstream effector of EPAC signaling. We also show that EPACs promote melanoma growth by regulating redox homeostasis and mitochondrial ROS through activation of mechanistic target of rapamycin complex 1 (mTORC1) that stabilizes hypoxia-inducible factor 1-alpha (HIF-1α), a transcriptional activator of redox regulator TXNIP and glycolytic enzymes. Our data suggest that targeting mechanisms that melanoma cells employ to bypass EPAC dependency is a potential therapeutic approach.

**Graphical Abstract:** 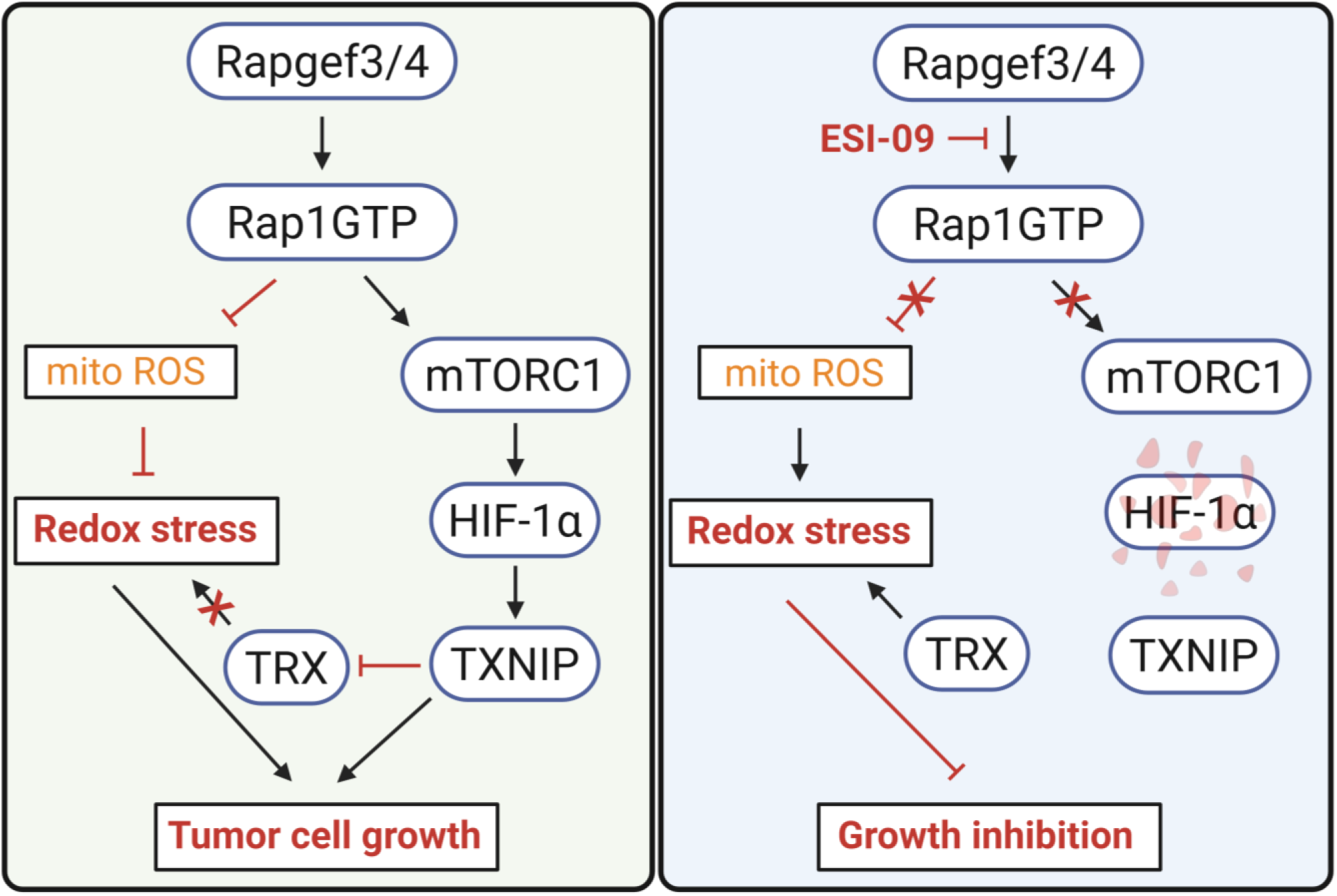

## INTRODUCTION

Cutaneous melanoma is a deadly neoplasm of skin melanocytes with high metastatic propensity. Despite advances in targeted therapy and immune checkpoint blockade, the limited effectiveness and durability of these treatments and development of resistance remain as major hurdles. Oxidative stress plays a critical role in metastatic progression of melanoma through reversible metabolic changes that enhance their capacity to withstand oxidative stress [1]. In a genetic model of mouse melanoma, antioxidant treatment was shown to promote metastases [2]. These reversible metabolic adaptations, which are critical for survival and growth, are induced by local tumor microenvironment and cellular redox homeostasis. However, the molecular mechanisms that drive these adaptations are not completely understood.

Exchange proteins activated by cyclic AMP (EPAC1/2) are RAP guanine nucleotide exchange factors [3]. EPACs have been implicated in the regulation of redox homeostasis in many normal cells and tissues and in immune modulation [4, 5]. In cancer cells, EPACs are known to regulate cell proliferation, migration and invasion [6]. We previously showed that EPAC signaling preferentially regulates the growth of primary melanoma cells by stimulating mTORC1 activity and loss of dependency on EPAC signaling correlates with melanoma progression [7, 8]. We also showed that inhibition of EPAC increased mitochondrial ROS production only in primary melanoma cells consistent with the role of EPAC-RAP1 signaling in maintaining mitochondrial integrity by suppressing mitochondrial ROS production and oxidative damage [5, 9]. The role of EPACs in melanomagenesis *in vivo* and the molecular mechanisms through which EPACs selectively promotes the growth of primary melanoma cells remain to be understood.

In this study, we show that expression of EPACs is induced early during transformation of melanocytes and EPAC activity is required for the growth of the transformed melanocytes. Employing *Braf/Pten* mouse melanoma model, we show that systemic inhibition of EPACs with ESI-09 or conditional deletion of *Epac1/2* in melanocytes delayed melanomagenesis and inhibited tumor growth. RNAseq analysis of patient-matched primary and lymph node-metastatic melanoma (LN-met) cells treated with ESI-09, revealed that TXNIP, a regulator of cellular oxidative stress, as a critical gene regulated by EPAC. Mechanistic studies revealed that EPAC signaling regulates TXNIP expression by stabilizing its transcriptional regulator HIF-1α, which also controls the expression of glycolytic enzymes. Analyses of TCGA data and quantitative immunohistochemical analysis (IHC) of a clinically annotated tumor tissue microarray (TMA) showed that low EPAC mRNA and RAP1-GTP (active RAP1) protein levels correlate with better disease (recurrence)-free survival of patients diagnosed with primary melanoma. These data suggest that EPACs are induced early in melanomagenesis and contribute to primary tumor growth by regulation of redox homeostasis, mitochondrial ROS and glycolytic metabolism through activation of mTORC1-HIF-1α-TXNIP signaling. We propose that while targeting EPAC signaling is an attractive strategy for prevention and interception of localized cutaneous melanoma, targeting the mechanisms that metastatic melanoma cells employ to bypass EPAC signaling is a novel treatment option for advanced melanoma.

## MATERIALS & METHODS

### Cell lines & chemicals

Primary and LN-met melanoma cell lines WM35, WM75, WM1552, WM1862C, WM115, WM266-4, WM239A, and WM165-1 were cultured in Tumor Specialized Medium (TSM) [10] with 2% heat-inactivated FBS and 1% penicillin and streptomycin (pen+strep). Metastatic melanoma cell lines (MRA2, MRA5, MRA6, MRA9, MRA12, and MRA13), gifted by Dr. Mark Albertini (UW-Madison), YUMM series cell lines, and *Epac*-null mouse tumor cell lines were cultured in DMEM with 10% FBS and pen+strep. Human melanocytes were isolated and cultured as described previously [11]. All cells were cultured in a humidified chamber at 37°C with 5% CO_2_. Cell lines were authenticated by short tandem repeat analysis by UW-TRIP lab. All Cell lines were routinely tested for mycoplasma contamination. Details of all the cell lines, chemicals, and antibodies used in this study are listed in Supplementary Table 1.

### Animals

*Braf/Pten (B6. Cg-Braf^tm1Mmcm^ Pten^tm1Hwu^ Tg (Tyr-Cre/ERT2)13Bos/BosJ*) mice were obtained from the Jackson Laboratory. The C57BL6/J mice with *Epac1/2* floxed alleles were a gift from Dr. Cheng of University of Texas-Houston [12]. *Epac^fl/fl^* mice were crossed with *Braf^V600E/+^/Pten^fl/fl^*(*BPT*) mice [13] to generate double heterozygous *BPT/Epac1*^+/*fl*^*/Epac2^+/fl^,* double homozygous (*BPT/Epac1^fl/fl^/Epac2^fl/fl^)*, *Epac1^fl/fl^* (*BPT/Epac1^fl/fl^)* and *Epac2^fl/fl^ (BPT/Epac2^fl/fl^)* as described previously. The genotype was confirmed by Transnetyx (Memphis, TN). Mice were kept in a controlled environment with free access to water and a standard laboratory diet. For experiments, both flanks of 3-4 weeks male and female mice *Braf^V600E/+^/Pten^fl/fl^*(*BPT* control), *Epac1^fl/fl^, Epac2^fl/fl^, Epac1^+/fl^/Epac2^+/fl^,* and *Epac1 ^fl/fl^/Epac2^fl/fl^* were shaved and 2 µL of 5 mM 4HT was applied topically on both flanks for two days to generate *BPT, Epac1 and Epac2 or Epac1/2* conditional knockout mice referred to hereafter as *BPT, Epac1^+/-^/Epac2^+/-^, Epac1^-/-^/Epac2^-/-^, Epac1^-/-^*, and *Epac2^-/-^,* respectively. Mice were monitored for palpable tumors. Tumor dimensions were measured with a digital caliper twice weekly until they reached the endpoint, i. e., 20 mm in any direction. Mice were euthanized by CO_2_ and tumors were excised. Half of each resected tumor was flash frozen and stored at -80°C for downstream biochemical analysis, and the remaining half was fixed in a 4% paraformaldehyde solution for histology. Some tumor tissues were processed for establishing cell lines and used for genotyping. All animal experiments were approved by Institutional Animal Care and Use Committees of both University of Wisconsin-Madison and Madison VA Hospitals.

For human tumor xenograft model, 4–6-week-old NSG (*NOD.Cg-Prkdc^scid^ Il2rg^tm1Wjl^/SzJ*; The Jackson Laboratory-USA) mice were subcutaneously inoculated with human primary WM115 (4x10^6^) and metastatic MRA6 melanoma cells (2x10^6^) in 100 μL PBS into the shaved flank. After tumor cell injection, mice were monitored for palpable tumors, and when the tumor size reached 100 mm^3^, mice were randomized into four treatment groups: control group (PBS for 7 days/week); ESI-09 group (7 days/week, 25mg/kg); EPACi+BRAFi group (ESI-09 7 days/week, 25mg/kg; vemurafenib 5 days/week, 10mg/kg) and BRAFi+MEKi group (vemurafenib 5 days/week, 10 mg/kg; Selumetinib 3 days/week, 10mg/kg). All compounds were administered intraperitoneally for three weeks. After three weeks of treatment, mice were euthanized, and excised tumors were weighed.

For treatment with ESI-09, 4HT was applied to 3–4-week-old *Braf/Pten* mice and randomized animals on day 21 received either vehicle (8:1:1 PBS/Tween-80/ethanol) or ESI-09 (25mg/kg) intraperitoneally for three weeks. Tumor measurements were recorded twice a week. Mice were euthanized once tumors reached the endpoint and tumors were excised.

### Tissue microarrays

We employed an in-house constructed human primary melanoma tissue microarray (TMA) described previously [14]. A TMA containing both primary and metastatic melanomas was acquired from US Biomax (ME483). The TMA slides were deparaffinized in xylene and were then rehydrated using a standard-graded ethanol series. Antigen retrieval (AR) was performed using microwave treatment and an AR buffer with pH values of 6.0 and 9.0. The TMA slides were stained using DAPI and the Opal seven-color IHC Kit and following combination of primary antibodies and fluorophores: S100 (Opal 520), Ki67 (Opal 570), RAP1-GTP (Opal 650) and RAP1-GAP (Opal 690). The slides were stained by repeating staining cycles consecutively, with microwave treatments between each cycle and the process was completed by staining with DAPI. The slides were imaged using Vectra 2.0 automated slide scanner using a scanning protocol based on core size and layout, as well as an acquisition of spectral library. For each slide, an 8-bit image cube from each of the TMA tissue cores was acquired and the fluorescence signals were analyzed using Nuance and Inform software 2.2.1 (PerkinElmer). Inform advanced image analysis software was used to segment tissues (melanoma versus non-melanoma) to analyze the signal intensity. Then, the target signals were quantified after unmixing the spectral curves. Continuous signal intensity (percentage positivity in selected tissue category area) was generated for each identified tissue/cell type. Quantitation for Ki67, RAP1GAP, and RAP1-GTP was performed in S100-positive cells. Pearson’s correlation coefficient was used to determine the correlation between Ki67, RAP1-GAP, and RAP1-GTP based on percentage positive staining.

### RNAseq analysis

Patient-matched primary and LN-met melanoma cell lines (WM115, WM266, WM239 and WM165) were treated with ESI-09 at 6 and 12 h. Cells were lysed, RNA was extracted and the RNA was submitted to UW-Madison Gene Expression Center for RNAseq. Raw FASTQ analysis data was uploaded to the Rosalind RNA seq analysis platform, HyperScale architecture developed by ROSALIND, Inc. San Diego, CA) (https://www.rosalind.bio). Briefly, reads were trimmed using cutadapt [15]. Quality scores were assessed using FastQC [16]. Reads were aligned to the Homo sapiens genome build GRCh38 using STAR [17]. Individual sample reads were quantified using HTseq [18] and normalized via Relative Log Expression (RLE) using the DESeq2 R library[19]. Read distribution percentages, violin plots, identity heatmaps, and sample MDS plots were generated as part of the QC step using RSeQC [20]. DEseq2 was also used to calculate fold changes and p-values and perform optional covariate correction. Clustering of genes for the final heatmap of differentially expressed genes was done using the PAM (Partitioning Around Medoids) method using the fpc R library [21]. The hypergeometric distribution was used to analyze the enrichment of pathways, Gene ontology, domain structure, and other ontologies. The topGO R library[22] was used to determine local similarities and dependencies between GO terms to perform Elim pruning correction. MSIgDB database source was referenced for enrichment analysis[23, 24]. Enrichment was calculated relative to a set of background genes relevant for the experiment. RNA seq data were analyzed for differential gene expression (DEG) between primary and LN met cell lines. The raw RNAseq data was deposited to a Gene Expression Omnibus (GEO) data set maintained by the National Institute of Health.

### TCGA data analysis

Skin Cutaneous Melanoma (TCGA, Firehose Legacy) was accessed on January 25, 2024 through cBioPortal [25, 26], includes mRNA expression and clinical outcomes information of 479 melanoma patients. This cohort comprised 100 individuals diagnosed with primary melanoma, with comprehensive clinical follow-up for 86 patients. We analyzed the relationship between the expression of *RAPGEF3* and *RAPGEF4* mRNA and disease-free survival (DFS) in patients diagnosed with primary melanoma based on high (≥ median) and low (≤ median) levels of both *RAPGEF3* and *RAPGEF4* to assess the cumulative effect of these two genes on DFS [27].

### Immunoblotting

Cells were washed, scraped, and lysed on the plate using RIPA buffer comprising a Halt protease inhibitor cocktail. The lysates were then sonicated and centrifuged for 30 min at 4°C to separate the supernatant. Protein concentration in the supernatant was estimated using the BCA protein assay, and 30-50 μg of total protein was resolved on SDS-PAGE gels and transferred onto a PVDF membrane. Membranes were blocked with a quick blocker for 10 min, followed by overnight incubation with respective primary antibodies at 4°C. After washing, membranes were incubated with respective horseradish peroxidase labeled secondary antibody for 60 min and followed by enhanced chemiluminescent (ECL) detection reagents and imaged using a LICOR Bio Odyssey FX imager.

### Cell growth assay

Cells were seeded in flat bottomed 96 well plate at 5000 cells/well and allowed to attach at 37°C overnight. The next day, cells were treated with various compounds (ESI-09, ESI-05, NAC, GSH, rotenone and MitoQ) for 72 h or five days. At the endpoint, 10 μL of 3-[4,5-dimethylthiazol-2-yl]-2,5diphenyltetrazolium bromide (MTT agent) was added to the wells, plates were incubated for 4 h at 37°C, followed by 100 μL of solubilization buffer. After overnight incubation at 37°C absorbance at 570 nm was measured using a Biotek Synergy H1 Multi-Mode Plate Reader.

In addition to MTT assays, we also performed cell counting with a Countess Automated Cell Counter (Applied Biosystems) in experiments where we treated primary melanoma cell lines WM115 and WM1862 with the EPAC inhibitor ESI-09 and transfected with EPAC siRNA. We found that neither ESI-09 treatment nor EPAC KD significantly changed cell numbers after 72 hours. These observations allowed us to interpret the MTT data as proxy for cell growth.

### qRT-PCR

Total RNA was extracted using RNeasy plus universal RNA isolation kit according to the manufacturer’s instructions. RNA was reverse transcribed using High-capacity RNA-cDNA reverse transcription Kit on a Mini Amp Plus Thermal Cycler (Thermo Fisher, Waltham, MA), and qRT-PCR was performed using TaqMan Fast Advanced master mix on a CFX96 Real-Time System (Bio-Rad, Hercules, CA). The following TaqMan gene expression assays were used in our study: GAPDH, RAPGEF3, RAPGEF4, TXNIP, ARRDC4, and HIF-1A. The reaction conditions were 50°C for 2 min (1 cycle), 95°C for 20 sec (1 cycle), 95°C for 3 sec and 60°C for 30 sec (40 cycles). The 2^-ΔΔCt^ method was used to calculate relative mRNA expression from Ct values. GAPDH gene was used as an internal reference.

### EPAC siRNA transfection

All knockdown experiments were performed using silencer select siRNAs (siEPAC1 and siEPAC2) purchased from Thermo Fisher. WM115 and WM165-1 cells were seeded at 1x10^6^ cells/well in triplicates in 6-well plates 24 hours before transfection. Cells were transfected with siEPAC1 or siEPAC2 and pooled siRNAs at 0.1 nM concentration each using Lipofectamine RNAiMax transfection reagent according to manufacturer’s instructions. After 48h, cells were harvested for western blotting analysis or cell counting by Countess cell counter (Invitrogen, Waltham, MA).

### HIF-1*α* (p402A/p564A) over-expression

Primary melanoma cells (WM115) were seeded at 1x10^6^ cells/well in triplicates in 6-well plates 24 hours before transfection. Cells were transfected with empty vector (EV) or degradation resistant HIF-1α (p402A/p564A) plasmid using Lipofectamine RNAiMax transfection reagent according to manufacturer’s instructions. After 48h, cells were harvested and plated into 96 well plates for cell growth assessment using MTT assay. The remaining cells were lysed and used for western blot analysis of HIF-1 α and TXNIP levels.

### TXNIP over-expression

Lentiviruses were produced in HEK293T cells using pMD2, psPAX2, and TXNIP expression plasmid using Lipofectamine 3000. The lentiviruses were collected at 24, 48 and 72h of transfection, titters were estimated, pooled, aliquoted and stored at -80°C after 0.22 μM filtration. For TXNIP overexpression (TXNIP-OE), 2x10^6^ primary melanoma cells (WM115 and WM1862) were seeded in a 6 cm culture dish and incubated with empty vector or TXNIP expression plasmid lentiviral particles in polybrene (8 μg/ml) containing medium four times with 6-8h intervals. After 48 hours of transduction, cells were maintained in G418 medium for selection. The cells were trypsinized 72h after transduction and plated into 96 well plates for cell growth assessment using MTT assay. The remaining cells were lysed and TXNIP overexpression was confirmed by western blot analysis.

### Establishment of mouse tumor cell lines

*Epac1^+/-^/Epac2^+/-^* and *Epac1^-/-^/Epac2^-/-^*mouse tumor cell lines were derived from tumors that reached endpoint volume (20 mm). The *Epac1^+/-^/Epac2^+/-^* (double heterozygous) and *Epac1^-/-^ /Epac2^-/-^* (double homozygous) tumors were dissected with a sterile scalpel blade, and a small piece was removed from the core of the tumor and transferred to the media and minced into small pieces and transferred into a 6 cm culture dish and cultured in DMEM with 10% FBS and pen+strep.

### Statistical analysis

All quantitative data are presented as mean ± standard deviation (SD) of multiple replicates as appropriate. Data shown for cell culture experiments are representative of 2-3 independent experiments. A p-value of less than 0.05 was considered statistically significant. To determine statistical significance, we performed an Unpaired t-test, a multiple unpaired t-test, and a One-way ANOVA with Tukey’s multiple comparison test. Data was analyzed and plotted with GraphPad prism 10 software (GraphPad Inc., USA).

## RESULTS

### EPAC-RAP1 signaling is correlated with aggressiveness of cutaneous primary melanoma

To investigate the prognostic value of EPAC1/2 (RAPGEF3/4) expression, we queried TCGA Firehose Legacy SKCM dataset, which consists of tumor mRNA expression and patient follow up data for a cohort of 86 patients diagnosed with primary melanoma. In this cohort, high expression (≥ median) of EPAC1/2 mRNA was associated with worse disease-free survival (HR=3.22, p= 0.0015) (**Fig. 1A**). To assess whether EPAC activity (i. e., RAP1-GTP levels) is associated with melanoma cell proliferation, we performed multiplexed quantitative IHC of a commercially available melanoma TMA (consisting of 32 Stage II and III primary lesions and 16 lymph node metastases) for RAP1-GTP protein and the proliferation marker Ki67 (**Fig. 1B**, **C**). We assessed the expression of RAP1-GTP and Ki67 in S100A (a melanoma marker) positive cells. There was a significant and positive correlation between RAP1-GTP protein and Ki67 positivity in primary lesions (**Fig. 1D**) but not in lymph node metastases (**Fig. 1E**) consistent with our previous finding on the loss of dependency of metastatic melanoma cells on EPAC-RAP1 signaling. We performed a similar analysis using an in-house built clinically annotated human primary melanoma TMA. Two representative lesions on the TMA with high and low RAP1-GTP and the corresponding Ki67 staining are shown in **Fig. 1F**. Quantitative analyses of RAP1-GTP and Ki67 in S100A-positive cells showed a weak but significant correlation between RAP1-GTP levels and proliferative cells **(Fig. S1A, B)**. The details of patients included in the in-house and commercially acquired TMA are shown in **Fig. S1C-E**. Kaplan-Meier analysis showed a significant difference in recurrence - free survival (DFS) between patients with high and low RAP1-GTP melanomas (log-rank test; p=0.0289), i. e., patients with high RAP1-GTP melanoma had higher probability of metastatic recurrence (HR: 5.50, 95% CI: 1.19-25.38 **(Fig. 1G).** These data suggest that EPAC-RAP1 signaling correlates with the proliferative capacity and is associated with poor prognosis.

**Figure. 1:**
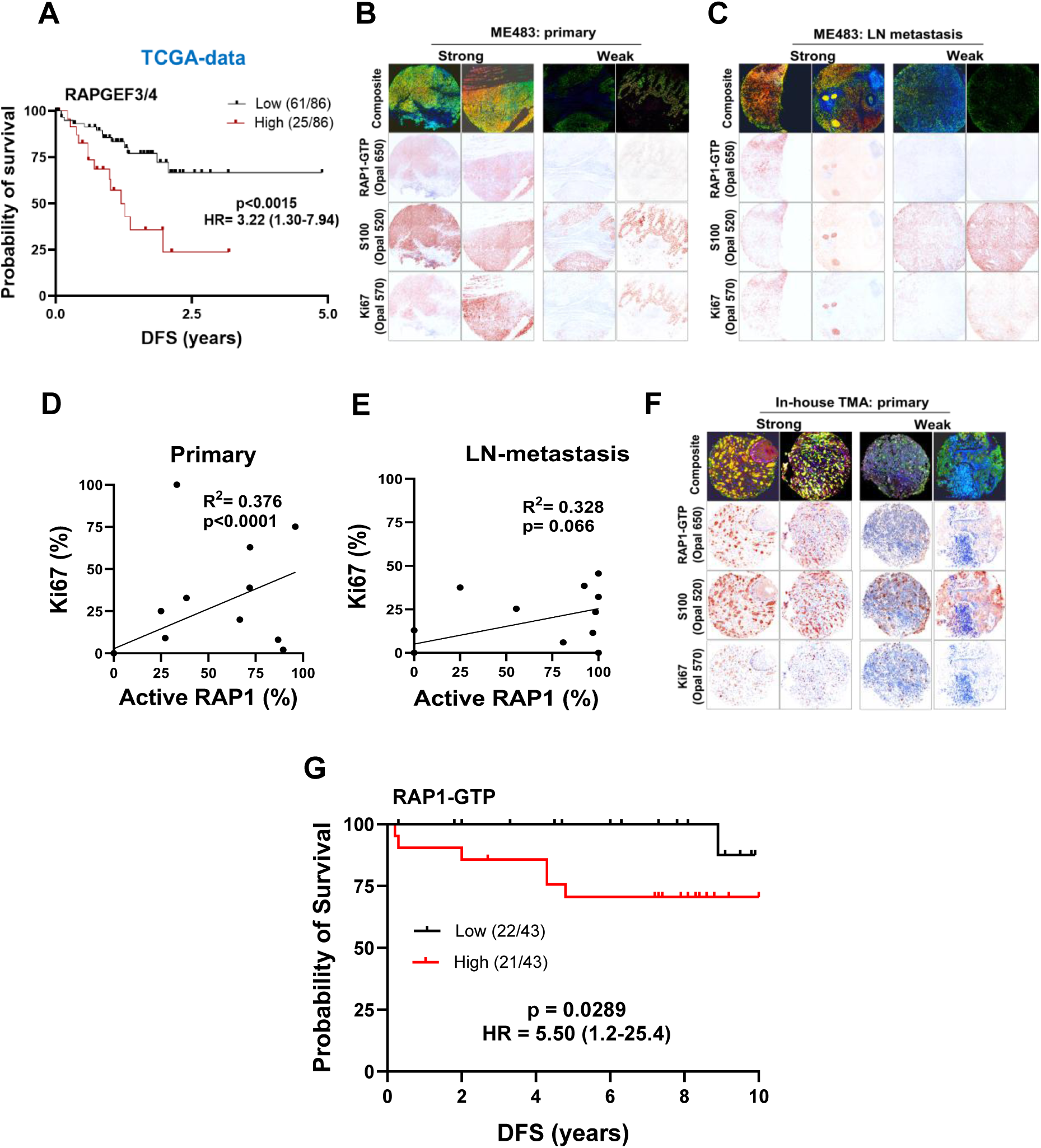
EPAC1/2 expression and activity correlate with primary melanoma tumor aggressiveness. **A.** Kaplan-Meier survival analysis of EPAC1/2 (RAPGEF3/4) gene expression and disease-free survival (DFS in years) from melanoma TCGA data set. Patients were stratified based on median (high or low) expression of EPAC1/2 mRNA. **B** and **C**. IHC analysis of a commercially available TMA consisting of primary and metastatic melanoma tumors. TMA was stained for RAP1-GTP, melanoma marker S100A, and proliferation marker Ki67 using Opal multiplex immunofluorescence kit. **D** and **E.** Linear regression analysis of RAP1-GTP protein and Ki67 in primary and metastatic melanoma tumors. **F.** Staining of an in-house TMA consisting of primary melanoma patient tumors for RAP1-GTP, S100A and Ki67 using Opal multiplex immunofluorescence kit. **G.** Kaplan-Meier survival analysis of data from primary melanoma patient TMA stained for RAP1-GTP expression. Patients were stratified based on median (high or low) expression of RAP1-GTP protein.

### Expression of EPAC1/2 is upregulated by PI3K-AKT signaling in melanocytes and EPAC2 activity is required for the growth of transformed melanocytes

To test whether EPAC signaling is activated and plays a role during malignant transformation of melanocytes, we transformed cultured neonatal human melanocytes (NHM) with lentiviruses for oncogenic BRAF (BRAF^V600E^) and/or short hairpin RNA (shRNA) for PTEN (phosphatase and tensin homolog). Expression of BRAF^V600E^ and knockdown (KD) of PTEN markedly upregulated EPAC1 and EPAC2 mRNA and proteins (**Fig. 2A, B**). Interestingly, PTEN-KD alone induced higher expression of both EPAC1 and EPAC2 compared to expression of BRAF^V600E^. In support for the role of PI3K-AKT signaling in EPAC1/2 upregulation, treatment with inhibitors of PI3K (KY12420) or AKT (KRX0401) abolished the EPAC1/2 expression in the transformed melanocytes (**Fig. S2A**). Intriguingly, combination of BRAF^V600E^ expression and PTEN-KD further increased EPAC2 expression but dampened EPAC1 expression (**Fig. 2B**). Upregulation of EPACs is also accompanied by increased activity as seen by the increased RAP1-GTP levels (**Fig. 2C**). To test whether upregulated EPAC activity is required for the survival and growth of transformed melanocytes, we treated the BRAF^V600E^/PTEN-KD transformed cells with EPAC1/2 inhibitor ESI-09 or EPAC2-selective inhibitor ESI-05 [28, 29]. As shown in **Fig. 2D**, treatment with either inhibitor decreased the cell growth. In addition, we transfected BRAF^v600E^/shPTEN transformed melanocytes with siEPAC1 or siEPAC2 and/or siEPAC1/2. siRNA-mediated knockdown (KD) of EPAC2 alone resulted in inhibition of growth in these transformed melanocytes, similar to the combined treatment of siEPAC1/2 (**Fig. S2B)**, showing that EPAC2 is the predominant drivers of the growth of these transformed melanocytes.

**Figure. 2:**
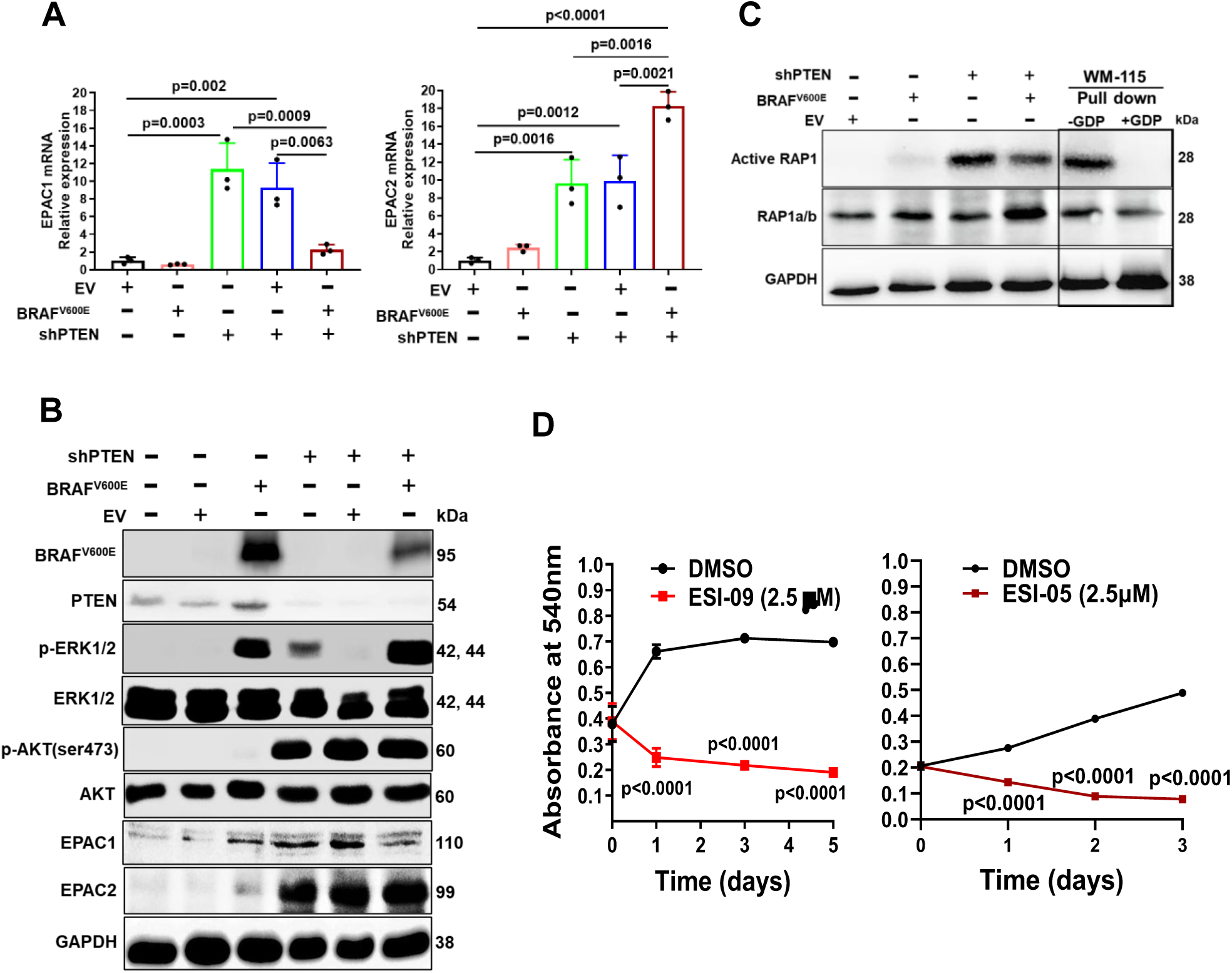
EPACs are activated during malignant transformation and promotes the growth of transformed melanocytes. **A.** EPAC1 and EPAC2 mRNA expression levels in transformed neonatal human epidermal melanocytes. **B.** Western blot analysis of EPAC1 and EPAC2 expression in normal and transformed melanocytes. Cells were cultured and transduced with lentiviruses for BRAF^V600E^ and for shPTEN for 72h. On day 6, cell lysates were prepared and subjected to SDS-PAGE and probed for indicated proteins. **C.** Western blot analysis of total and RAP1-GTP in transform melanocytes. RAP1-GTP pull-down from WM115 human primary melanoma cells was used as a positive control and GDP treated WM-115 cells used as a negative control. GAPDH was used as loading control. **D.** Effect of EPAC1/2 inhibition by ESI-09 or EPAC2 inhibition by ESI-05 on survival of human transform melanocytes cells. Melanocytes were cultured and then transduced with lentivirus with shPTEN, BRAF^V600E^ for 5days. On day 6, cells plated in 96-well plates were treated with ESI-09 (2.5 µM) or ESI-05 (2.5 µM) for 5 days; drug was replenished every 48 hours. Data normalized to DMSO control are shown as mean ± SD of 6 replicates and analyzed using Student’s t-test and p-values are shown for each data point.

Since expression of BRAF^V600E^ in melanocytes is known to induce oncogene-induced senescence [30], we asked whether EPAC activity induced by PTEN-KD contributes to the senescence escape mechanisms activated by PI3K-AKT signaling. We treated cells transduced with BRAF^V600E^ alone and both BRAF^V600E^/shPTEN with ESI-09 or ESI-05 and stained them for senescence associated β-galactosidase (SA-β-gal) activity (**Fig. S2C**). Treatment with either EPAC1/2 or EPAC2 inhibitor significantly increased the percentage of SA-β-gal-positive cells (**Fig. S2D**). Western blot analysis showed that inhibition of EPAC resulted in downregulation of cyclins E1, A2, and B1 and cell cycle dependent kinases CDK4 and CDK2, and concomitant upregulation of cell cycle inhibitors p16, p21 and p27 (**Fig. S2E**). Overexpression of EPAC2, on the other hand, dampened senescence and promoted the growth of transformed melanocytes (**Fig. S2F**). These data suggest that upregulation of EPACs is an early event in melanomagenesis and EPAC activity contributes to the suppression of oncogene-induced senescence and promotes the growth of malignant melanocytes.

### Systemic inhibition of EPAC1/2 activity delays melanoma tumor growth *in vivo*

To test the role of EPACs in melanomagenesis, we initiated melanoma development in immunocompetent *Braf/Pten* (*BPT*) mice by topical application 4HT (4-hydroxytamoxifen) on the shaved flanks. On day 21 of post-4HT application, we randomized the mice and administered vehicle or EPACi (ESI-09; 25mg/kg) intraperitoneally for three weeks. All animals in the vehicle treated group reached the endpoint of tumor burden (when animals had to be euthanized) by 83 days whereas EPACi delayed the tumor growth and the mice survived up to170 days (**Fig. 3A, B** and **Fig. S3)**. Kaplan-Meier survival analysis showed that EPACi significantly improved the survival of *BPT* mice (median survival 111 days, HR: 0.3829; CI: 0.096-1.519; p ≤ 0.05) compared to vehicle-treated mice (**Fig. 3C, D**). These results show that EPAC signaling promotes the growth of *Braf/Pten* driven melanoma.

**Figure. 3:**
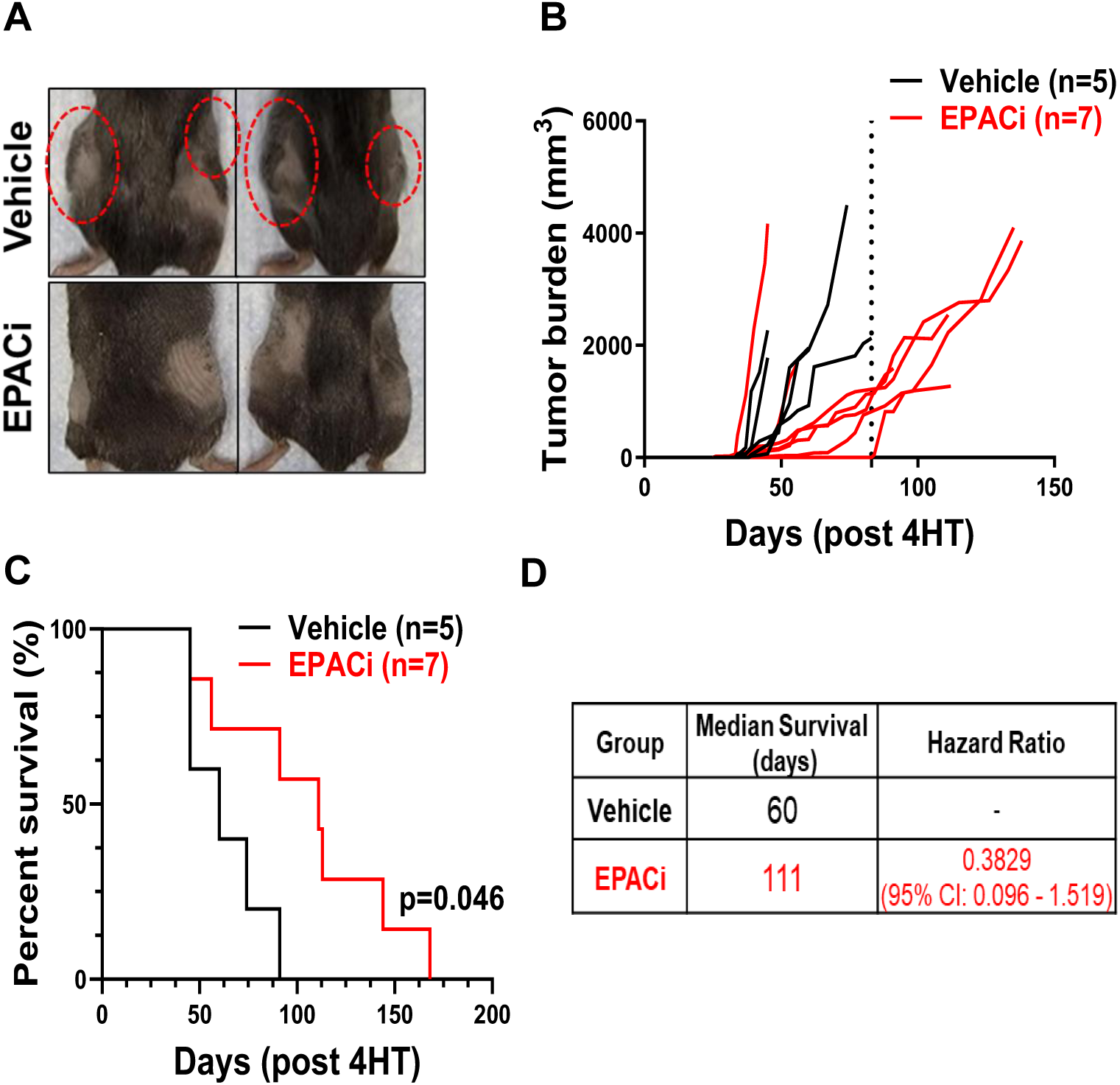
Systemic inhibition of EPAC activity delayed melanomagenesis and tumor growth in *Braf/Pten (BPT)* mouse melanoma model. **A.** Representative images of flanks of mice show the difference in tumor development between vehicle and EPACi treated *BPT* mice at week five after 4HT application. The red circles represent tumors on the flanks of mice. **B.** Tumor burden in control EPACi treated mice. Each line in the plot represents an individual mouse. Total tumor burden on both flanks is shown. **C.** Kaplan-Meier survival analysis of vehicle and ESI-09 treated mice. Log-rank (Mantel-Cox) test was performed for survival trends between experimental groups. **D.** Table showing median survival and hazard ratio values between experimental groups. The median survival, hazard ratio, and CI were determined from the Kaplan-Meier survival curve and log-rank (Mantel-Cox) test.

### Genetic deletion of *Epac1* and *Epac2* reduces melanoma tumor growth

We developed mice with conditional deletion of *Epac1* and *Epac2* genes individually or in combination, in the immunocompetent *BPT* mouse model. On shaved flanks of 3–4-week-old *BPT* mice or *BPT*-*Epac1^fl/fl^* (*Epac1^-/-^*) and -*Epac2^fl/fl^* (*Epac2^-/-^*), and *BPT*-*Epac1^+/fl^/Epac2^+/fl^* (*Epac1^+/-^ /Epac2^+/-^*) and *BPT*-*Epac1^fl/fl^/Epac2^fl/fl^*(*Epac1^-/^*^-^*/Epac2^-/^*^-^) mice, we applied 4HT to initiate melanomagenesis. Presence of the knockout alleles of *Epac1/2* in the tumor tissues was confirmed by genotyping (**Fig. S4**). Five weeks after 4HT application, *BPT* control mice showed palpable tumors that grew rapidly whereas *Epac1^-/-^* mice exhibited a decrease in tumor incidence (p ≤ 0.0004) and *Epac2^-/-^* mice exhibited longer tumor latency (p ≤ 0.002) but there was no significant difference in tumor burden or overall survival compared to the control mice (**Fig. S5**).

However, when measurable tumors appeared in *BPT* mice around 5 weeks, *Epac1/2* double heterozygous (*Epac1^+/-^/Epac2^+/-^)* and double homozygous (*Epac1^-/^*^-^*/Epac2^-/^*^-^) mice had significantly smaller or no palpable tumors (**Fig. S6A** and **Fig. 4A**). The tumor latency for *BPT* mice was 29.6±1.6 days compared to 63.1±7.8 in double heterozygous mice and 52.9±7.2 days in double homozygous mice, although only the increase in latency was significant (p < 0.04) in the double heterozygous mice (**Fig. 4B**). *BPT* mice developed tumors on both flanks showing a 97% incidence with maximum tumor burden observed by 94 days. Double heterozygous mice exhibited a significantly lower tumor incidence of 60.4% (p ≤ 0.0003) and maximum tumor burden by 254 days; double homozygous mice showed a tumor incidence of 90% with a maximum tumor burden by180 days. (**Fig. 4C, D**). Kaplan-Meier survival analysis showed that both double heterozygous and double homozygous mice had better survival compared to *BPT* control. The median survival for *BPT* mice was 56 days whereas the median survival of double heterozygous mice was 95 days (HR 0.3278; CI: 0.111-0.964; p ≤ 0.003) and 123 days for double homozygous mice (HR 0.2444; CI: 0.074-0.797; p ≤ 0.0001) (**Fig. 4E, F**). These data show a role for *Epac1/2* in melanomagenesis and tumor growth. Histochemical (hematoxylin and eosin staining) and IHC staining (with melanoma marker S100A) of excised tumors confirmed the melanocytic origin of the tumors **(Fig. S6B)**.

**Figure. 4:**
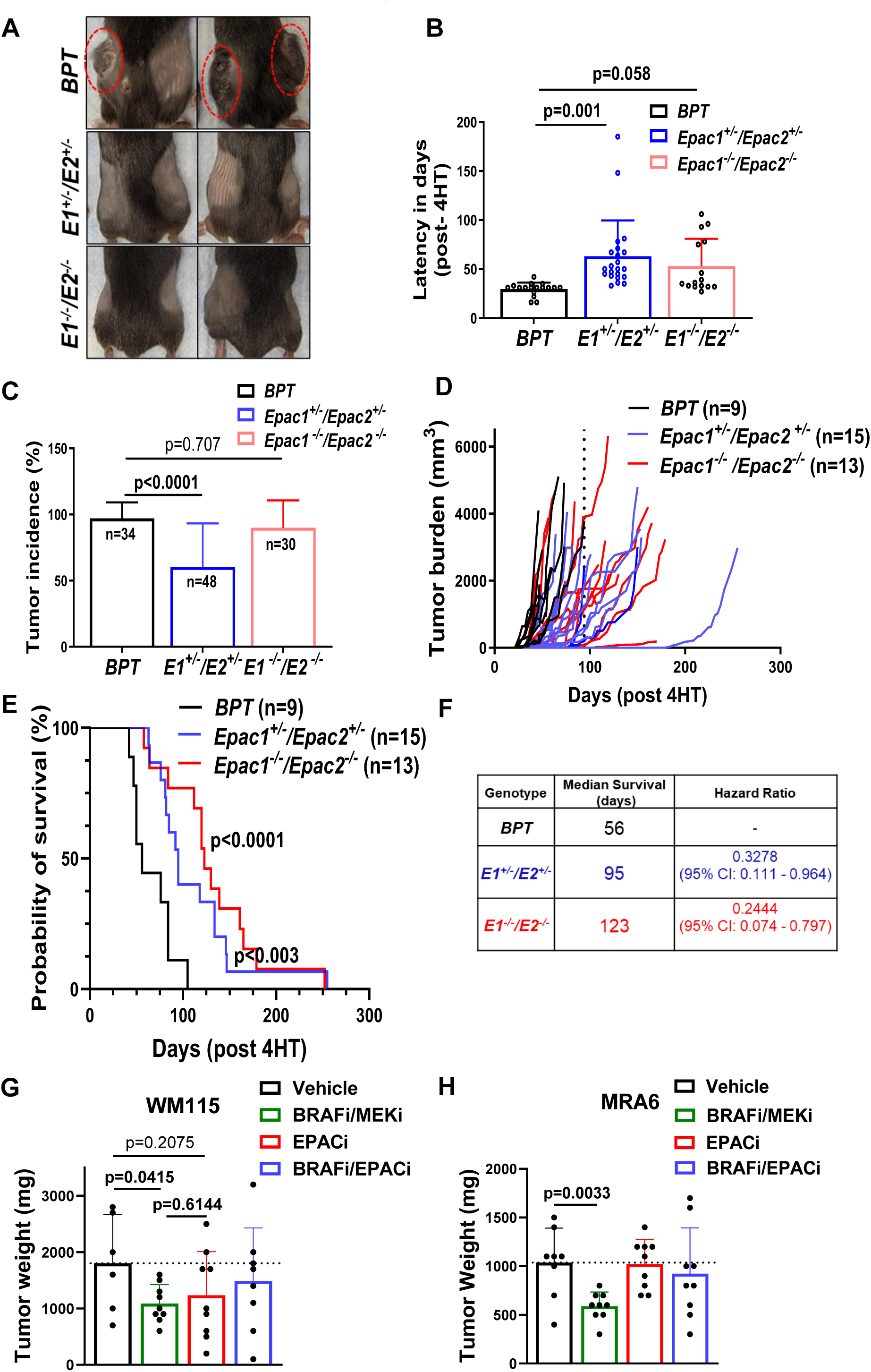
Deletion of *Epac1/2* reduced tumor growth and improved overall survival in mice. **A.** Representative images of flanks of *BPT*, *Epac1/2-*double heterozygous and double homozygous mice showing differences in tumor development at week five after 4HT application. The red circles represent tumors on the flanks of *BPT* mice. **B.** Melanoma tumor latency (days) in *BPT, Epac1/2-*double heterozygous and double homozygous mice. Data are shown as mean ± SD and analyzed using One-way ANOVA using Tukey multiple comparison test. **C.** Comparison of tumor incidence between *BPT, Epac1/2-*double heterozygous and double homozygous mice. n = number of tumors induced in each group. Two tumors were induced per mouse (each flank one tumor). Tumor incidence is presented as percent incidence. Data are shown as mean ± SD and analyzed using One-way ANOVA using Tukey multiple comparison test. **D.** Tumor burden in *BPT, Epac1/2-*double heterozygous and double homozygous mice. Each line in the plot represents tumor burden in individual mouse. **E.** Kaplan-Meier survival analysis of *BPT, Epac1/2-*double heterozygous and double homozygous mice. Log-rank (Mantel-Cox) test was performed for survival trends between experimental groups. **F.** Table showing median survival and hazard ratio values between experimental groups. The median survival, hazard ratio, and CI were determined from the Kaplan-Meier survival curve and log-rank (Mantel-Cox) test. **G** and **H.** Tumor weights from NSG mice xenografted with primary WM115 and metastatic MRA6 melanoma cells treated with vehicle or indicated agents. Data were shown as mean ± SD of tumor weight and analyzed using Unpaired t-test.

EPACs have been shown to modulate immune response [4, 31]. To test whether immune response plays inhibition of melanoma tumor growth by loss of *Epac1/2*, we assessed the frequency/abundance of tumor resident CD8^+^ and CD4^+^ T cells. Immunofluorescence staining of tumor sections showed a higher frequency of CD8^+^ cells in *Epac1/2* double heterozygous (19.03%) and double homozygous (9%) tumors compared to *BPT* tumors (5.8%) (**Fig. S7A, B**). All tumor sections stained with anti-CD4^+^ antibody showed weak or no immunofluorescence signal. These data suggest that EPAC1/2 expression in melanoma cells is associated with reduced tumor resident CD8^+^ cells.

To further test the relationship between immune cells and the role of EPAC1/2 in melanoma growth, we inoculated human primary WM115 and metastatic MRA6 melanoma cells (both from mutant BRAF melanomas) subcutaneously into the flanks of immunodeficient NSG mice. When the tumors reached 100 mm^3^, we randomized the mice into four groups and treated them with vehicle or BRAFi (Vemurafenib) and MEKi (Selumetinib), or EPACi (ESI-09) alone or EPACi with BRAFi for three weeks (**Fig. S7C).** On Day 22, we excised tumors and measured the tumor weight **(Fig. S7D, E**). As expected, BRAFi/MEKi treatment significantly inhibited the growth of both primary and metastatic BRAF mutant tumors (WM115, p=0.0415; MRA6, p=0.003). Interestingly, EPACi treatment of mice with either WM115 or MRA6 did not affect the tumor growth. Moreover, in both WM115 and MRA6 injected mice, treatment with EPACi and BRAFi resulted in no significant decrease in tumor weight compared to vehicle (**Fig. 4G, H**). These data show that the inhibitory effect of EPACi on primary melanoma growth is less pronounced in immune deficient mice and suggest suppression of immune cell mediated mechanisms also contribute to EPAC-dependent growth promotion of melanoma *in vivo*.

### Gene Expression Changes induced by EPAC inhibition

To understand the mechanism of action of EPAC in primary melanoma, and the differential dependency of primary and metastatic melanoma cells on EPAC, we performed RNAseq analysis of matched primary WM115 and LN-met melanoma cell lines WM266-4, WM239A and WM165-1 treated with ESI-09. Differentially expressed gene (DEG) analysis showed that in WM115 cells treated for 6h with ESI-09, only two genes, *TXNIP* and *ARRDC4,* were significantly downregulated (fold change ≥ 2; p ≤ 0.05), whereas only *TXNIP* was downregulated in WM165-1 cells. After 12h treatment of WM115 cells with ES-09, 288 genes were downregulated and 332 were upregulated (fold change ≥ 2; p ≤ 0.05). Interestingly, in WM165-1 cells, even after 12h treatment with ESI-09 there were no additional up- or downregulated genes apart from *TXNIP* (**Fig. 5A**). DEGs in primary melanoma after 12h ESI-09 treatment is shown in the volcano plot, and the heat map shows the top 25 up- and downregulated genes (**Fig. 5B, C**). Pathway enrichment analysis using MSIgDB dataset (https://www.gsea-msigdb.org) showed that most significantly altered top 4 Hallmark pathways are: E2F_TARGETS, G2M_CHECKPOINT, GLYCOLYSIS and TGF_BETA_SIGNALING (**Fig. 5D** and **Fig. S8A**).

**Figure. 5:**
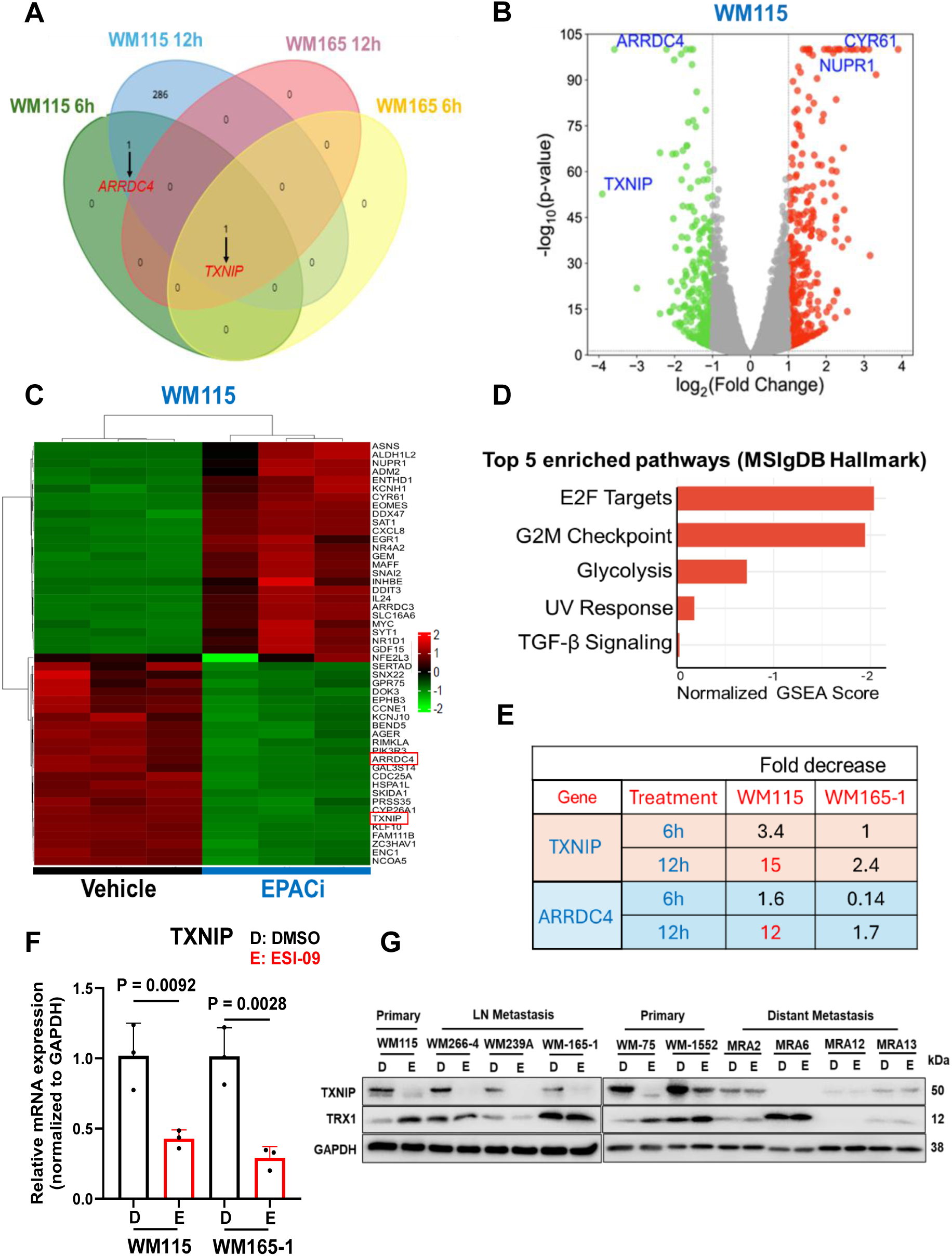
Gene expression analysis of primary and LN-met melanoma cell lines treated with EPAC1/2 inhibitor ESI-09. **A.** Venn diagram showing number of genes that are downregulated (fold change ≥ 2; p ≤ 0.05) in primary and LN-met cells treated with ESI-09 for 6 and 12h. **B.** Volcano plot for changes in gene expression in primary melanoma WM115 cell line treated with ESI-09 for 12h [Red: upregulated genes; Green: downregulated genes. Data were plotted using SRplot [77]]. **C.** Heatmap of upregulated and downregulated genes in WM115 primary melanoma cell line treated with ESI-09 for 12h [fold change ≥ 2; p ≤ 0.05; Hiplot [78]]. **D.** Gene enrichment analysis (MSIgDB hallmark gene sets) showing top 5 enriched pathways. **E.** Table showing fold decrease in TXNIP and ARRDC4 genes in primary and LN-met cells treated with ESI-09 for 6 and 12h. **F.** qRT-PCR validation of TXNIP expression in WM115 and WM165-1 cell lines treated with ESI-09 for 12h. GAPDH is used as an internal control. Fold change in TXNIP expression determined by using 2^-ΔΔCt^ method. Data are presented as mean ± SD, analyzed using One-way ANOVA with Tukey multiple comparison test and p-values are shown. **G.** Western blot validation of downregulation of TXNIP expression using a panel of primary and metastatic melanoma cell lines treated with ESI-09 for 24h (**D:** DMSO; **E:** ESI-=09).

TXNIP, a member of α-arrestin protein family, binds to the reduced forms of thioredoxins - cytosolic TRX1 and mitochondrialTRX2- and is an important regulator of redox homeostasis [32–34]. RNAseq analysis showed a large time-dependent decrease (3.4-fold at 6h and 15-fold at 12h) in TXNIP mRNA in ESI-09 treated WM115 cells but a smaller effect (2.5-fold decrease at 12h) in WM165-1 (**Fig. 5E**). qRT-PCR and western blot analysis of a panel of primary and metastatic melanoma cell lines treated with DMSO or ESI-09 showed more abundant expression of TXNIP mRNA and protein in primary melanoma cell lines (**Fig. S8B** and **Fig. 5F**) and confirmed the decrease in TXNIP mRNA and protein expression in ESI-09 treated primary and LN-melanoma cell lines but not in distant metastatic melanoma cell lines (**Fig. S8C** and **Fig. 5F)**. Consistent with its role in regulating TRX levels and function, downregulation of TXNIP by ESI-09 resulted in upregulation of TRX1 protein in all primary (but not metastatic) melanoma cell lines (**Fig. 5G)**. ARRDC4 (Arrestin domain containing protein-4), also a member of the arrestin family, regulates glucose transporter system [35]. Although qRT-PCR analysis confirmed downregulation of ARRDC4 mRNA by ESI-09 in all primary cell lines tested, there was no difference in basal expression of ARRDC4 mRNA between primary and metastatic cell lines, and we did not observe any marked decrease in protein levels in different primary melanoma cell lines treated with ESI-09 (**Fig. S8D-F**).

### TXNIP is a downstream effector of EPAC signaling

To confirm the role of EPACs in regulation of TXNIP, we transfected primary WM115 and WM1862 and LN-met WM165-1 melanoma cells with EPAC1/2 siRNA. As previously reported [7], siRNA-mediated knockdown (KD) of EPAC1/2 or EPAC2 alone resulted in growth inhibition (based on cell counting using automated Cell Counter) in both primary melanoma cell lines but not LN-met cells. Western blot analysis showed that EPAC-KD is accompanied by a decrease in TXNIP protein **(Fig. 6A, B)**. We also tested TXNIP expression in mouse melanoma cell lines established from *Epac1/2* double heterozygous and double homozygous tumors. Genotyping of these cell lines confirmed the knockout of *Epac* genes **(Fig. S9A)**. Western blot analysis of these tumor cell lines showed BRAF^V600E^ expression, loss of PTEN and reduced TXNIP expression (**Fig. S9B** and **Fig. 6C**).

**Figure. 6:**
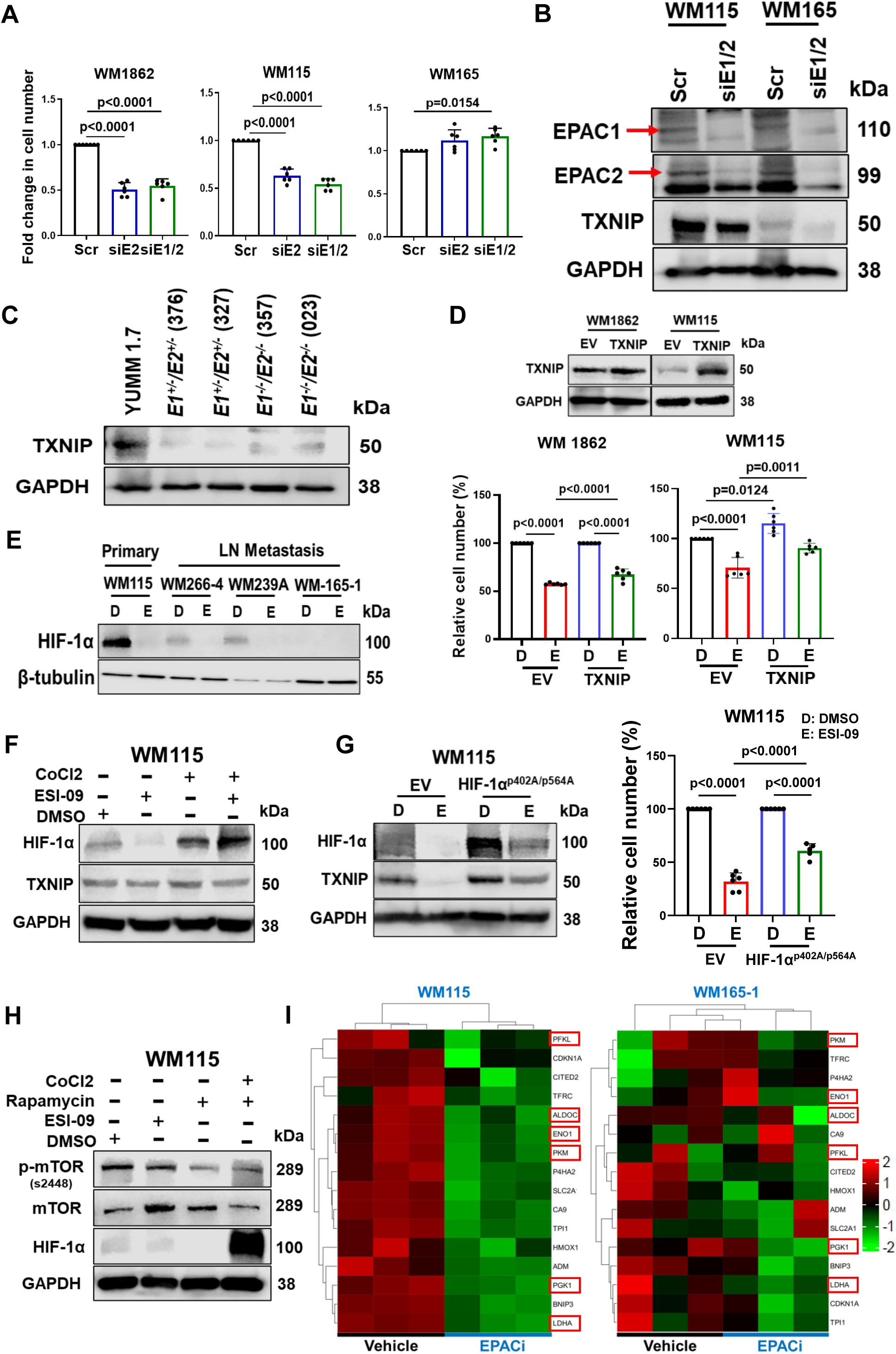
EPAC regulates TXNIP expression through mTORC1-HIF-1*α* signaling. **A.** Primary (WM115 & WM1862) and LN-met (WM165-1) melanoma cells were transfected with control scrambled (NC) or EPAC1/2 siRNA. Cell numbers were estimated 72 h post-transfection. Data are presented as mean ± SD, analyzed using One-way ANOVA with Tukey multiple comparison test and p-values are shown. **B.** Western blot analysis of EPAC1, EPAC2, and TXNIP expression in EPAC1/2siRNA and scrambled control-transfected primary (WM115) and LN-met (WM165) melanoma cells. **C.** Western blot analysis of TXNIP in tumor cell lines derived from *Epac* null mice tumors. **D.** *Top:* Western blot validation of TXNIP overexpression in two different primary (WM1862 & WM115) melanoma cell lines transduced with TXNIP lentivirus for 48h. *Bottom:* Effect of TXNIP overexpression on growth inhibition by ESI-09 in WM1862 and WM115 primary melanoma cells (**D**: DMSO; **E**: ESI-09). Data shown are relative cell numbers and presented as mean ± SD, analyzed using One-way ANOVA with Tukey multiple comparison test and p-values are shown. **E**. Western blot analysis of HIF-1α expression in patient matched primary and LN-met melanoma cells treated with ESI-09 for 6h (**D**: DMSO; **E**: ESI-09). **F.** Western blot analysis of HIF-1α and TXNIP expression changes in WM115 primary melanoma cells treated with ESI-09 or CoCl_2_ alone and in combination for 6h. **G.** *Left panel:* Western blot analysis of HIF-1α overexpression and TXNIP levels in DMSO and ESI-09 treated primary (WM115) melanoma cells transfected with degradation resistant mutant HIF-1α (p402A/p564A) plasmid for 24h. *Right panel:* Effect of HIF-1α (p402A/p564A) overexpression on growth inhibition by ESI-09 (**D**: DMSO; **E**: ESI-09). Data shown are relative cell numbers and presented as mean ± SD, analyzed using One-way ANOVA with Tukey multiple comparison test and p-values are shown. **H.** Western blot analysis of mTOR, phospho-mTOR (ser-2448), and HIF-1α expression changes in WM115 primary melanoma cells treated with ESI-09 or rapamycin alone or rapamycin with CoCl_2_ for 6h. GAPDH is used as a loading control. **I.** Heatmap of HIF-1 target genes [MSigDB Human Gene Set: SEMENZA_HIF1_TARGETS [79]] in primary and LN-met melanoma cell lines treated with ESI-09 for 12h. Downregulated glycolytic enzymes in primary melanoma are highlighted in red boxes.

To test the role of TXNIP in the growth inhibitory effect of EPACi, we asked whether overexpression of TXNIP (TXNIP-OE) can relieve the effect of EPACi. Data in **Fig. 6D** show that TXNIP-OE in primary melanoma cell lines WM1862 and WM115 dampened the growth inhibition caused by EPACi showing that TXNIP is a downstream effector of EPAC signaling in primary melanoma.

### TXNIP expression is regulated by EPAC-mTORC1-HIF-1*α* axis

To understand the mechanisms involved in regulation of TXNIP by EPAC signaling, we investigated the expression changes in transcription factors known to regulate TXNIP. These include MONDOA (Max-like protein X interacting protein), FOXO1A (Forkhead box O1A), FOXO3 (Forkhead box O3), HIF1α (Hypoxia inducible factor 1α) and NRF2 (Nuclear factor erythroid 2-related factor 2) [36–40]. Examination of RNAseq data did not reveal any significant changes in the expression of MONDOA, FOXO1A, FOXO3, and NRF2 mRNA, and western blot validation of protein in ESI-09 treated primary and LN-met cells (**Fig. S10A, B**). Interestingly, although HIF1A mRNA showed a significant increase only in WM115 cells, qRT-PCR analysis did not validate this increase (**Fig. S10C**). Western blot analysis, on the other hand, showed that ESI-09 treatment abolished the HIF-1α protein expression as early as 6h in primary and LN-met cell lines (**Fig. 6E)** with sustained downregulation up to 72h **(Fig. S10D**).

To test whether EPAC inhibition results in downregulation of HIF-1α protein levels by affecting its accumulation, we treated primary WM115 cells with ESI-09 alone or cobalt chloride (CoCl_2_; 100 μM), which stabilizes HIF-1α protein by inhibiting hydroxylation by prolyl hydroxylases, or combination of ESI-09 and CoCl_2_ for 6h. Western blot analysis showed that addition of CoCl_2_ resulted in HIF-1α accumulation and prevented downregulation by ESI-09 treatment. More importantly, accumulation of HIF-1α by CoCl_2_ alone resulted in increased TXNIP levels and prevented its downregulation by ESI-09 (**Fig. 6F & Fig. S10E)**. To test whether overexpression of degradation-resistant HIF-1α protein rescues TXNIP levels in ESI-09 treated cells, primary melanoma cells (WM115) were transfected with empty vector or plasmid encoding a degradation-resistant HIF-1α (HIF-1α p402A/p564A) protein. As shown in Fig. 6G (*left panel*), overexpression of the mutant HIF-1α alone increased TXNIP levels and rescued it from downregulation by ESI-09 treatment. Over expression of the mutant HIF-1α also relieved primary melanoma cells from growth inhibition by ESI-09 treatment (**Fig. 6G**, *right panel*). These data suggest that EPAC signaling regulates TXNIP expression by allowing maintenance of high levels of HIF-1α protein.

We previously showed that EPAC-RAP1 signaling regulates mTORC1 activity[7]. mTORC1 signaling is known to upregulate HIF-1α activity by both increasing its transcription and protein stabilization [41, 42]. In agreement with the critical role of mTORC1 signaling in regulation of HIF-1α, we found that treatment of WM115 primary melanoma cells with rapamycin also resulted in a marked decrease in HIF-1α protein **(Fig. 6H)** that could be rescued by addition of CoCl_2_.

HIF-1α is also an important regulator of transcription of metabolic enzyme genes [43]. We, therefore, queried our RNAseq dataset for changes in the expression of HIF-1α regulated metabolic enzymes (SEMENZA_HIF1_TARGETS in MSIgDB). **Fig. 6I** shows expression of several glycolytic enzymes such as phosphofructokinase liver type (PFKL), aldolase C (ALDOC), pyruvate kinase M1/2 (PKM), phosphoglycerate kinase-1 (PGK1), enolase-1 (ENO1), and lactate dehydrogenase A (LDHA) downregulated in ESI-09 treated primary WM115 cells suggesting that EPAC signaling regulates primary melanoma cell metabolism by stabilizing HIF-1α.

### EPAC signaling regulates TXNIP expression independent of redox homeostasis

We previously showed that EPAC regulates mitochondrial ROS production preferentially in primary melanoma cells[7]. TXNIP shuttles from the nucleus to mitochondria in response to oxidative stress and regulates mitochondrial oxidative stress [44]. We treated primary and metastatic melanoma cells with MitoQ (150 nM), the triphenyl phosphonium (TPP) conjugated antioxidant ubiquinone, which acts as a mitochondrial antioxidant in normal cells. In cancer cells, MitoQ, on the other hand, has been shown to increase mitochondrial oxidative stress and reduce cell survival including that of melanoma cells [45]. Like the treatment with ESI-09, MitoQ treatment reduced the growth of primary cells and metastatic melanoma cells are less sensitive to MitoQ. Interestingly, treatment with both ESI-09 and MitoQ further suppressed the growth of primary melanoma cells compared to the treatment with either agent (**Fig. 7A)**. Additionally, treatment with rotenone (1 μM), an inhibitor of mitochondrial complex I electron transport chain[46], markedly inhibited the growth of primary melanoma cells but not LN and distant metastatic cells **(Fig. S11A)**. These data support the notion that EPAC promotes the growth of primary melanoma cells by dampening mitochondrial oxidative stress.

**Figure. 7:**
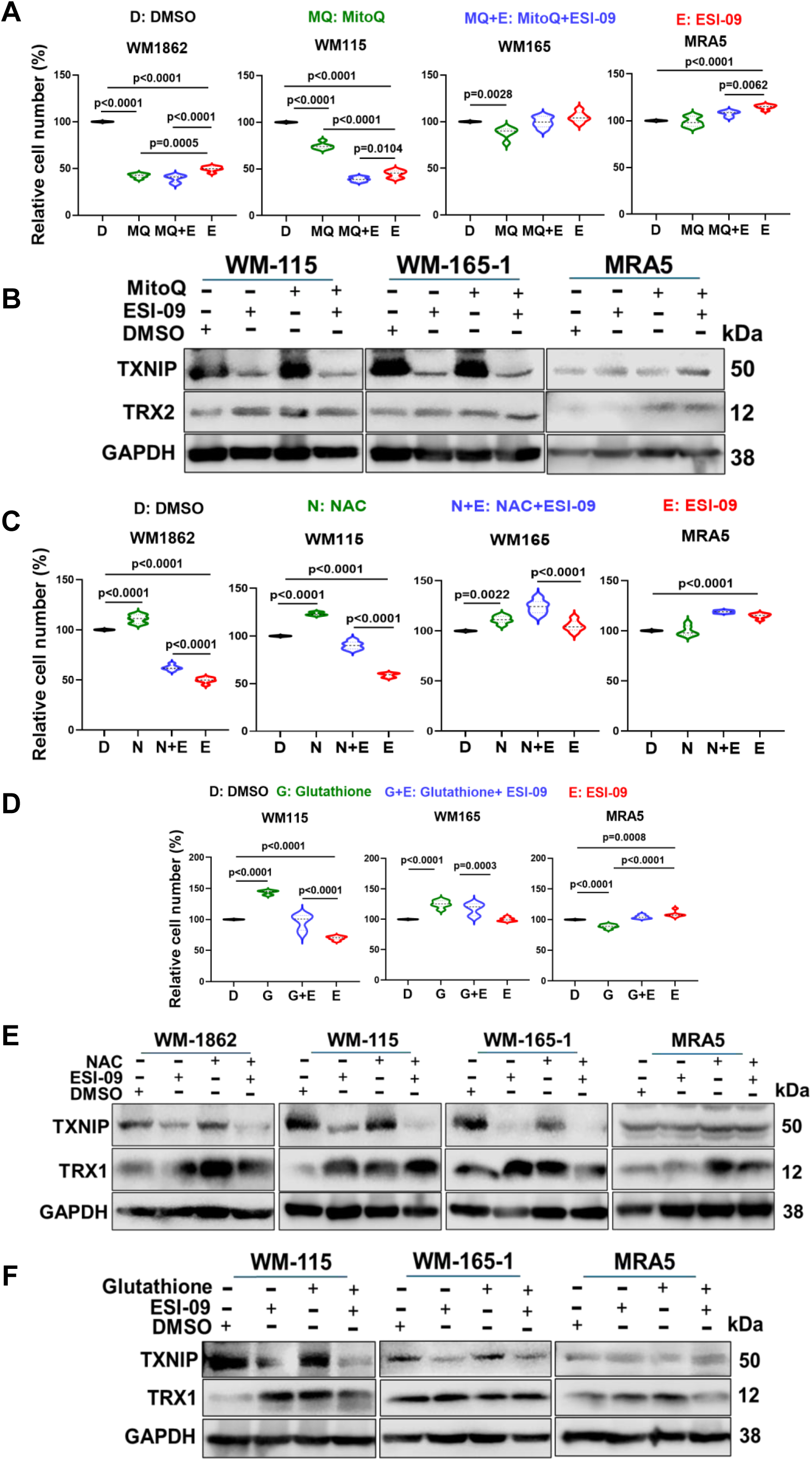
Effect of antioxidants on the growth of primary and metastatic melanoma cell lines and relationship to TXNIP expression: **A.** Primary (WM1862, WM115), LN-met (WM165-1), and distant met (MRA5) cells were plated in six replicates in a 96-well plate (5000 cells/well) and treated with 2.5 µM ESI-09 or 150 nM MitoQ alone or in combination for 72h. Cell numbers were estimated using MTT assay. Data are presented as mean ± SD, analyzed using One-way ANOVA with Tukey multiple comparison test and p-values are shown. **B**. Western blot analysis of TXNIP and TRX1 from WM115, WM165-1, MRA5 cells treated with ESI-09 or MitoQ alone or in combination for 24h. **C.** Primary and metastatic cells were plated as in **A** and treated with 2.5 µM ESI-09 or 1 mM NAC alone or in combination for 72h. Cell numbers were estimated using MTT assay. Data are presented as mean ± SD, analyzed using One-way ANOVA with Tukey multiple comparison test and p-values are shown. **D.** Primary and metastatic cells were plated as in **A** and treated with 2.5 µM ESI-09 or 0.6 mM GSH alone or in combination for 72h. Cell numbers were estimated using MTT assay. Data are presented as mean ± SD, analyzed using One-way ANOVA with Tukey multiple comparison test and p-values are shown. **E.** Western blot analysis of TXNIP and TRX1 from WM1862, WM115, WM165-1, MRA5 cells treated with ESI-09 or NAC alone or in combination for 24h. **F**. Western blot analysis of TXNIP and TRX1 from WM115, WM165-1, MRA5 cells treated with ESI-09 or GSH alone or in combination for 24h.

Treatment with MitoQ did not affect TXNIP levels or the inhibition of TXNIP by ESI-09 treatment. However, there was an increase in the mitochondrial TRX2 (presumably the oxidized form) in response to treatment with ESI-09 or MitoQ alone or in combination (**Fig. 7B**). These data show that growth inhibitory effect of MitoQ on primary melanoma cells is independent of TXNIP and suggest that EPAC signaling promotes melanoma cell growth through both TXNIP-dependent and -independent mechanisms.

EPACs are known to maintain low oxidative stress levels in different disease conditions [5, 9, 47]. To test ESI-09 treatment induces oxidative stress in melanoma cells, ESI-09 or antioxidant GSH treated primary (WM115, WM1862), LN-met (WM165-1) and distant metastatic (MRA5) cells were incubated with cell permeable dye CellROX green (binds to DNA upon oxidation and exhibits green fluorescence). ESI-09 treated WM115 and WM1862 cells showed more fluorescence signal compared to the control or GSH treated cells, indicating increased ROS production. As expected, we did not observe any changes in fluorescence signal in metastatic cells. These data support differential role for EPAC in maintaining redox homeostasis in primary and metastatic melanoma cell (**Fig. S11B, C**).

To further investigate the role of EPAC in regulation of melanoma cell oxidative stress, we treated primary and metastatic cells with antioxidants N-acetyl cysteine (NAC) and glutathione (GSH) alone or in combination with ESI-09. Addition of NAC (1 mM) or GSH (0.6 mM) in the medium promoted the growth of primary and LN-met cells but not distant metastatic melanoma cells. Treatment with a combination of ESI-09 and NAC or GSH suppressed the growth stimulatory effects of both antioxidants (**Fig. 7C, D**) suggesting that EPAC plays a role in promoting the growth of primary melanoma cells by maintaining low oxidative stress and this dependency on EPAC is lost during melanoma progression. Treatment with NAC or GSH did not affect TXNIP levels or downregulation of TXNIP by ESI-09. Following NAC or GSH treatment, the levels of cytosolic TRX1, presumably the reduced form, were significantly elevated in both primary and metastatic melanoma cells independent of TXNIP levels (**Fig. 7E**, **F**) suggesting that antioxidant response to NAC or GSH is mainly mediated by TRX1, and this response is progressively diminished as melanoma progresses.

### EPAC signaling is not involved in phenotypic switching of primary melanoma cells

Malignant melanoma exhibits phenotypic plasticity with a dynamic switch between proliferative and invasive states [48]. These phenotypes are often represented by high MITF and E-cadherin expression in proliferative cells and high AXL and N-cadherin expression in invasive cell. To investigate whether ESI-09 induced growth suppression is a manifestation of phenotypic switch from proliferative to invasive state of primary melanoma cells, we analyzed changes in the expression of markers of phenotypic switch, and the invasive and migratory behavior of patient-matched primary (WM115, WM75) and LN-met cells (WM165-1, WM239A) and a panel of unrelated primary (WM1862, WM1552C) and metastatic (MRA7) cell treated with ESI-09. EPAC inhibition did not result in any consistent changes in the expression of these phenotypic markers across all primary and metastatic melanoma cells tested (**Fig. S12A**). Next, we assessed whether inhibition of melanoma cell growth by ESI-09 is accompanied by phenotypic switch, i. e., an increase in their invasive and migratory behavior using Matrigel and wound healing assays. EPAC inhibition resulted in no significant change in invasion (**Fig. S12B and C**) and migratory behavior (**Fig. S12D-F**) of melanoma cells suggesting that EPAC inhibition does not cause phenotypic switch.

Based on our data, we propose a model **(Fig. 8)** for the mechanisms of action of EPAC in promoting the growth of primary melanoma cells. These mechanisms include a) suppression of mitochondrial ROS production, and b) activation of mTORC1 signaling, which stabilizes HIF-1α to enhance transcription of glycolytic enzymes and TXNIP, a regulator of the key redox homeostasis protein TRX. Inhibition of EPAC signaling, therefore, inhibits cell growth due to cascading effects that result in increased cellular oxidative stress, decreased glycolytic activity and inhibition of cell cycle progression.

**Figure. 8:**
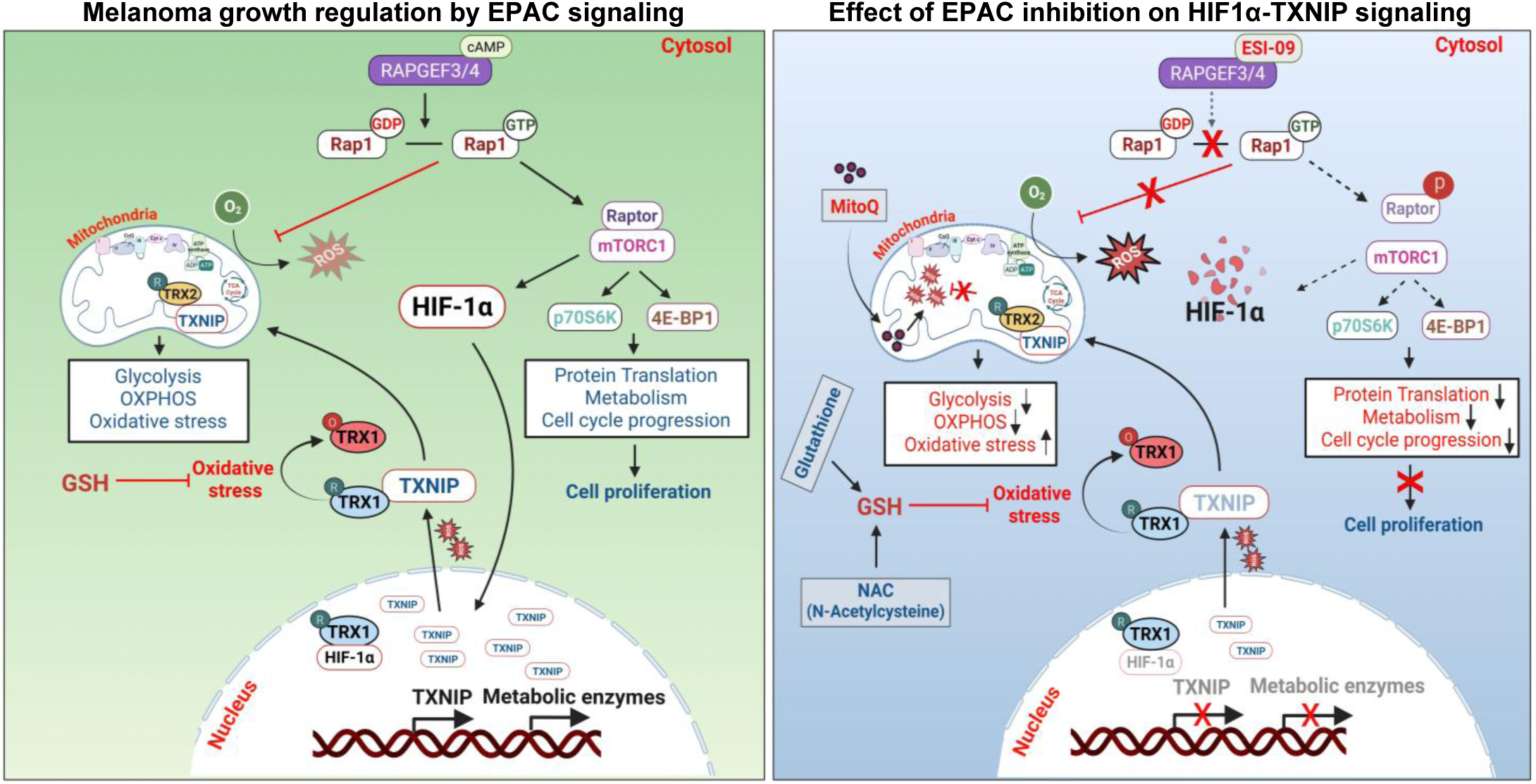
Proposed mechanism of action of RAPGEF3/4-RAP1GTP signaling in melanoma. The binding of cAMP (left panel) to RAPGEF3/4 (EPAC1/2) activates small G-protein RAP1 by the exchange of GDP with GTP. Activation of RAP1 promotes the activity of mTORC1 signaling. RAP1-GTP inhibits the production of mitochondrial reactive oxygen species (ROS). mTORC1 activity stabilizes HIF-1α protein and its nuclear translocation leading to the transcription of TXNIP mRNA and several glycolytic enzymes. Under oxidative stress, TXNIP relocates to cytosol and mitochondria, inhibiting the function of TRX1 and TRX2, respectively. The TXNIP-TRX1/2 interactions regulate metabolic activity, protein translation, cell cycle progression, supporting primary melanoma cell growth. Treatment with ESI-09 (right panel), a competitive inhibitor of cAMP binding to RAPGEFF3/4, inhibits guanine exchange factor activity of RAPGEFFs and RAP1 activation. Inactive RAP1 (RAP1GDP) promotes inhibitory phosphorylation of Raptor and inhibits mTORC1 activity [7]. Reduced mTORC1 activity results in the degradation of HIF-1α and downregulation of TXNIP expression. Under oxidative stress, lower TXNIP levels promote the activity of TRX1/2 leading to the inhibition of melanoma cell growth (BioRender).

## DISCUSSION

Mechanisms that promote metastatic progression of skin resident primary melanoma cells are not fully understood. In this study, we demonstrate EPAC-RAP1 signaling plays a critical role in promoting the growth of primary melanoma cells through multiple mechanisms that converge on regulation of cellular redox homeostasis, mitochondrial ROS production and glycolysis and propose that loss of this dependency on EPAC-RAP1 correlates with metastatic progression.

In this study, we show that melanocytes acquire this dependency on EPAC signaling during malignant transformation. In primary melanocytes, activation of PI3 kinase activity (by PTEN KD) alone resulted in upregulation of EPAC1/2 more dominantly than expression of BRAF^V600E^. Interestingly, in our study PTEN KD alone in primary melanocytes enhanced growth accompanied by the activation of EPAC2. Together with our observations on the requirement of EPAC2 (by treatment with EPAC2-selectibve inhibitor ESI-05 and overexpression of EPAC2) for growth and suppression of BRAF^V600E^-induced senescence, these data suggest that activation of EPACs, specifically EPAC2, is an early event in melanomagenesis. Previously, it was reported that EPAC1 protein, but not EPAC2, is overexpressed in metastatic melanoma tumors compared to primary melanoma. Based on increased migration of a highly metastatic cell line treated with EPAC-agonist 8-pMeOPT and reduced lung colonization of tail vein injected EPAC1-KD cells it was suggested that EPAC1 promotes migration [49, 50]. In this study, we employed both chemical inhibition and genetic deletion methods *in vivo* and multiple primary and patient-matched primary and metastatic melanoma cell lines *in vitro* to uncover the role of EPACs in melanomagenesis and growth of primary melanoma but not invasion or migration. In other cancers, EPACs were reported to either promote or inhibit cell proliferation, tumor growth and invasiveness, depending on the type of cancer [6, 51–56]. Our findings show that genetic deletion or systemic inhibition of EPAC prior to development of palpable tumors inhibited melanomagenesis in immunocompetent *Braf/Pten* models. On the other hand, EPAC inhibition did not affect tumor growth in immunocompromised NSG mice. Although this could suggest a role for immune system in EPAC-mediated melanoma growth promotion, we cannot rule out the possibility that the differences in the experimental design such as the dose, initiation and duration of treatment of the tumor bearing NSG mice with the EPAC inhibitor can explain these differences.

We identified thioredoxin interacting protein (TXNIP) as a gene regulated by EPAC signaling in melanoma cells. TXNIP is an important component of thioredoxin buffer system (TRX1/2, TXNIP, TRXR1/2, and NADPH) and is essential for cellular redox homeostasis [32–34]. This system is involved in the redox regulation of several proteins and transcription factors such as MYC, NF-κB, HIF-1α, and FOXOs [32–34]. TXNIP binds to reduced thioredoxin protein (TRX1/2) and inhibits its activity, thereby modulating cellular redox homeostasis [57]. TXNIP can act as either an oncogene or a tumor suppressor depending on the type and stage of cancer [58–61]. We found that TXNIP levels are decreased in metastatic lines compared to primary melanoma lines, and TRX1 protein levels are higher in metastatic cells compared to their matched primary cells. Meylan et al also found that TXNIP mRNA expression is significantly lower in Stage III and IV compared to healthy skin biopsies [62]. In a small set of 9 primary tumors, Knoll et al., reported that TXNIP expression and Breslow depth of invasion show an inverse correlation with low TXNIP levels at > 4 mm versus high levels at < 1 mm depth of invasion suggesting that TXNIP and components of its regulatory signaling axis can serve as novel prognostic markers for early stage metastasis [63]. Analysis of publicly available gene expression datasets GSE3189 and GSE46517 showed lower expression, respectively, of both EPAC2 and TXNIP in melanoma compared to melanocytic nevi [64] and higher TXNIP mRNA expression in nevi and a progressive decrease in TXNIP mRNA levels in primary to metastatic lesions [65]. Consistent with an inverse correlation between the levels of TXNIP and TRX, meta-analysis of public datasets GSE 15605, 38312 and 53223 revealed that TRX mRNA levels are higher in melanoma compared to melanocytes and nevi and it was suggested that TRX is a potential biomarker [66].

In this study, we report for the first time that EPAC signaling regulates TXNIP expression in melanoma cells. TXNIP is known to be regulated by different transcription factors [36–40]. We observed nearly complete downregulation of HIF-1α protein level as early as 6 hours of treatment with ESI-09 preceding a 15-fold downregulation of TXNIP mRNA seen at 12 hours. Moreover, overexpression of degradation-resistant HIF-1α relieved EPAC inhibition via increase in TXNIP levels.

Hypoxia-inducible factor-1α (HIF-1α) is a master regulator of cellular hypoxic stress, and cellular oxygen levels control the stability of HIF-1α protein [67]. Increased HIF-1α activity is well documented in several cancers, including melanoma [68]. Stabilization of HIF-1α is known to regulate tumor angiogenesis and metabolism, promoting tumor cell proliferation [43]. TXNIP expression was shown to be induced by HIF-1α in response to hypoxia in NSCLC and pancreatic cancers [37, 69]. However, there are no canonical HIF-1α binding sites in the *TXNIP* promoter region and there is no published evidence for binding of HIF-1α to the promoter region of *TXNIP* gene. These observations suggest that TXNIP expression is responsive to hypoxia and HIF-1 α regulates TXNIP by indirect mechanisms.

In previous studies, we showed that growth inhibition of primary melanoma cells by ESI-09 is accompanied by attenuated mTORC1 activity. mTORC1 is known to regulate the transcription and stability of HIF-1α protein [41, 42]. In mutant BRAF melanoma cells, although inhibition of BRAF^V600E^ by vemurafenib reduced HIF-1α protein, TXNIP and ARRDC4 were shown to be upregulated by binding of the alternative transcription factor MondoA to the promoters of these genes [70]. Therefore, it is possible that TXNIP expression in melanoma cells is regulated by EPAC-RAP1-mTORC1 signaling by both HIF-dependent and HIF-independent mechanisms. Other mechanisms for regulation of TXNIP and a role for TXNIP in proliferation of melanoma cells and EMT have been described [63, 71]. However, in this study we did not observe any changes in the expression of EMT markers or migratory and invasive capacity of patient-matched primary and metastatic cells treated with EPAC inhibitor that consistently downregulated TXNIP.

Oxidative stress is linked to tumorigenesis, tumor cell proliferation, and remodeling of tumor microenvironment [72, 73]. Thioredoxin system play a vital role in cancer cell antioxidant defense systems. Many cancer cells show elevated levels of GSH and TRX to combat high levels of ROS, which is linked to poor prognosis [74]. Tumor cells respond to oxidative stress by altering the activity of antioxidant transcription factors including HIF-1α essential for cellular redox homeostasis [72, 75]. TRX1 is known to stabilize HIF1α, facilitate its nuclear translocation and increase the transcription of hypoxia genes [76]. Oxidative stress is a major limiting factor for melanoma metastasis *in vivo*. Administration of antioxidants increased migratory and invasive capabilities of human melanoma cells *in vivo* by metabolic rewiring [1, 2]. In this and our previous study, we showed that inhibition of EPAC increases both cytoplasmic and mitochondrial ROS [7]. In kidney tubular epithelial cells, activation of EPAC-RAP1 signaling was shown to reduce the ROS production by preventing mitochondrial superoxide formation [9]. A role for EPAC-RAP1 signaling in suppressing mitochondrial ROS production in heart has also been documented [5], supporting a role for EPAC-RAP1 signaling in mitochondrial ROS homeostasis in melanoma.

In summary, we showed that EPAC regulates primary melanoma growth through multiple mechanisms involving i) RAP1GTP-mediated regulation of mitochondrial ROS production, ii) mTORC1 mediated protein translation, cellular metabolism and cell cycle progression, and iii) mTORC1-HIF1α mediated regulation of TXNIP/TRX, a critical regulator of redox homeostasis. Loss of dependency on EPAC for these cellular functions is associated with melanoma metastasis. We propose that identification of mechanisms that bypass EPAC dependency in metastatic melanoma will uncover novel targets for melanoma treatment independent of melanoma oncogenic drivers.

## Acknowledgements

This work was supported by VA Merit Awards I01BX004921 and 1I01CX002393, BLRD Research Career Scientist Award 1IK6BX006317 (to VS), UW Foundation’s Evan P. and Marion Helafer Professorship and the Department of Dermatology, University of Wisconsin-Madison; and NIH (R01CA261937), VA (I01BX005917, I01CX002210, VA Senior Research Career Scientist Award IK6BX006041), Dr. Frederic E. Mohs Skin Cancer Research Chair endowment).

## Author contribution

VS conceived the study and designed the research. SKT performed most of the experiments, interpreted the data, and wrote the manuscript draft. VS interpreted the data and wrote the manuscript. MKS carried out TCGA analysis, TMA immunostaining, RNA extraction, and submission for RNA-Seq analysis, as well as melanocyte experiments. SR carried out CD4/CD8 immunostaining, as well as invasion and migration assays. RR performed immunofluorescence staining, imaged and quantified CD8/CD4 stain data. SKT, SA, and SG performed cell line and mouse experiments. SKT, SG, and VS analyzed the mouse experimental data. SKT performed RNA-Seq data analysis. DDB analyzed histological data. MAN assisted with RNA-Seq data analysis. NA provided NSG mice for xenograft experiments. XC provided *Rapgef3^fl/fl^* and *Rapgef4^fl/fl^*mice and ESI-09. All the authors edited and approved the manuscript.

## Data availability

All data generated in this study are available in the article and the accompanying supporting data file. The RNA-Seq data have been deposited in the NCBI’s Gene Expression Omnibus (GEO) database (**GSE319527**). Complete Western blots are also provided in the supporting data.

## Study approval

Animal studies were approved by the IACUC of University of Wisconsin-Madison (protocol M006451) and VA (1594103-26). IRB approved human subject protocols (2016-0287 & 2006-006).

