## Supplementary Materials for "Thioredoxin interacting protein (TXNIP), a redox regulator, mediates the EPAC-RAP1 signaling dependency of primary melanoma"

#### Supplementary Methods:

##### Immunohistochemistry

Mouse tumor tissues *(BPT*, *Epac1/2*-heterozygous and homozygous) were Paraffin-embedded and sectioned by UW-TRIP lab. The first and last sections from each tumor tissue from given genotypes were H&E (Hematoxylin & Eosin) stained. For S100A immunostaining, tumor sections were dewaxed in xylene and hydrated in a decreasing gradient of alcohol. A quick melanin bleaching step was performed with 0.5% $H_2O_2$ in Tris-base (pH 10) for 15 min at 80$^o$ C (1). Antigen retrieval was carried out with Tris-EDTA buffer (pH - 9.0) in the microwave in three cycles (5 min each with 1 min interval). After PBS washes, nonspecific binding was minimized by protein blocker for 10 min at room temperature (RT). The sections were then incubated with primary antibody at 1:100 dilution for S100A in TBST with 5% BSA for 2 h 30 min at RT. After PBS washes, sections were incubated in HRP-labelled anti-rabbit secondary antibody for 30 min at RT. Staining was visualized by covering the sections in 3'-3' diaminobenzidine (DAB) buffer for 5 min, followed by counterstaining with hematoxylin. After subsequent dehydration in graded alcohol followed by xylene, sections were mounted, and images were captured under the light microscope (magnification: 200X).

##### Immunofluorescence

Tumor tissue sections *(BPT*, *Epac1/2*-heterozygous and homozygous) were dewaxed in xylene and hydrated in a decreasing gradient of alcohol. Antigen retrieval was carried out with citrate antigen unmasking buffer in the microwave in three cycles (5 min each with 1 min interval). After PBS washes, nonspecific binding was minimized by blocking sections with 5% donkey serum for 1h at RT. The sections were then incubated with primary antibody at a 1:200 dilution for CD8 and CD4 in TBST with 5% donkey serum overnight at 4°C. After PBS washes, sections were incubated in Alexa Fluor conjugated donkey anti-rat IgG secondary antibody for 60 min at RT. After PBS wash sections were mounted with antifade mounting media with DAPI. Images were captured using fluorescence microscope (EVOS core XL; magnification; 400X).

**Senescence-associated β-galactosidase assay**

Senescence-associated β-galactosidase (SA-β-gal) activity was assessed using the Senescence β-Galactosidase Staining Kit following the manufacturer's instructions. Briefly, melanocytes (3.5 × $10^3$ cells/well) were seeded into 24-well plates and transduced the following day with lentiviral vectors expressing empty vector control, $BRAF^{V600E}$, shPTEN, or the combination. After six days of transduction, cells were treated with ESI-09 or ESI-05 for 24 hours. On day 8, SA-β-gal staining was performed, and wells were imaged using an EVOS microscope. The proportion of SA-β-Gal–positive (blue) cells was quantified in ImageJ (Version 2.14.0/1.54f) by manual counting of stained cells relative to the total number of cells per field. Images were minimally processed using Adobe Photoshop for presentation.

**Detection of cellular ROS**

ROS levels in melanoma cells were analyzed by CellROX Green, a permeable green fluorescence dye, according to the manufacturer instructions. In brief, different primary (WM115, WM1862), LN-met (WM165) and distant metastatic (MRA5) melanoma cells were plated ($1X10^5$/well) in an 8-well glass bottom chamber slide and treated with ESI-09 or antioxidant GSH. After 24h, CellROX dye was added to cells at a final concentration of 5 μM, and cells were incubated for 30 min at37 °C. After incubation, cells were washed with PBS and fixed for 15min with 4% PFA followed by PBS washes. Finally, cells were mounted with antifade mounting media with DAPI and imaged by fluorescence microscope using 20X objective (EVOS core XL).

**Scratch Assay for Migration of Cells**

A panel of primary and metastatic melanoma cells ($1x10^5$/70 µL cells) were seeded into each chamber of the cell culture inserts placed in 24 well plates and incubated at 37 °C and 5% $CO_2$. After 24 hours, the inserts were removed to create a defined cell-free gap of 500 µm and the cells were treated with ESI-09 (2.5 µM) or DMSO (control). Cell migration was assessed at various time points, and the images were acquired at 0, 6, 24, and 48 h post-wounding using an EVOS core XL microscope using 10X objective. Quantification of wound area was performed using ImageJ.

**Matrigel Invasion Assay**

Cell invasion assay was performed using Matrigel Invasion Chambers (BD Bioscience) per the manufacturer's protocol. Briefly, 5 $x10^4$ cells in medium (without FBS) were seeded in Matrigel-coated trans wells in 24-well plates. Culture medium with 10% FBS was added to the lower chamber of the plate, which was further incubated at 37°C and 5% $CO_2$. After 48 h cells were stained with 1% crystal violet for 1 h. The cells that invaded the bottom of Matrigel membrane were imaged using an EVOS core XL microscope using 10X and 40X objectives. Quantification of cell numbers was performed using FIJI automated method (2).

**EPAC2 over-expression**

Lentiviruses were produced in HEK293T cells using pMD2, psPAX2, and EPAC2 expression plasmid (a generous gift from Dr. Stephen Yarwood of the University of Glasgow, UK) using Lipofectamine 3000. The lentiviral particle collection was performed at 24, 48, and 72h of transfection and stored at -80° C after 0.22 µM filtration. The titter was estimated according to the manufacturer's instructions. For EPAC2 overexpression $BRAF^{V600E}$/shPTEN transduced neonatal melanocytes ($2x10^6$ cells) were seeded and incubated with lentiviral particles packed with an empty vector, and EPAC2 overexpression plasmids in polybrene (8 μg/ml) containing medium four times with 6-8h intervals. After 48 hours of transient transduction, cells were maintained in G418 medium for selection. The cells were trypsinized 72h after transduction and plated into 96 well plates for MTT and senescence associated β-galactosidase assay. EPAC2 over expression was confirmed by western blot analysis.

**Supplementary Figures**

**Figure Legends:**

**Supplementary Figure. 1: EPAC1/2 expression and activity correlate with primary melanoma tumor aggressiveness. A.** Linear regression analysis of RAP1-GTP protein and Ki67 in in-house primary tumors. **B**. Linear regression analysis of RAP1-GTP protein and Ki67 in commercially available primary and metastatic melanoma tumors. **C** and **D.** Summary of clinicopathological information and univariant cox regression analysis data of patients from in-house TMA. Significant p-values are highlighted in red. **E.** Demographic information of patients from commercially available TMA (US Biomax-ME483).

**Supplementary Figure. 2: EPACs are activated during malignant transformation and promote the growth of transformed melanocytes. A.** Western blot analysis of EPAC1 and EPAC2 expression in transformed melanocytes treated with.PI3K and AKT inhibitors. Cultured neonatal melanocytes were transduced with $BRAF^{V600E}$ and for shPTEN for five days. The transduced melanocytes were treated with 1.5 μM KY12420 (PI3Ki) and 2 μM KRX0401 (AKTi) on day six for 16hr. Whole cell lysates were subjected to SDS-PAGE and probed for indicated proteins. **B.** *Left panel:* Western blot analysis of EPAC1 and EPAC2 knockdown (si*EPAC*) in $BRAF^{V600E}$/sh*PTEN* transformed melanocytes. *Right panel:* Effect of knockdown of EPAC2 and EPAC1/2 on the growth of transformed melanocytes. **C.** Representative microscopic image of transformed melanocytes treated with DMSO and EPAC1/2 inhibitor ESI-09. The red arrows show SA-β-gal positive cells transduced with oncogenic $BRAF^{V600E}$ or shPTEN and/or both $BRAF^{V600E}$/shPTEN. **D.** Cultured neonatal melanocytes were transduced with shPTEN, $BRAF^{V600E}$ lentiviruses for 7days. On day 8 cells were treated with 2.5μM ESI-09 and ESI-05 for 24h and β-galactosidase activity was measured. P-values are shown for comparisons between treatments. **E.** Cultured neonatal melanocytes were transduced with $BRAF^{V600E}$ and shPTEN lentiviruses for five days. The transduced melanocytes were treated with ESI-09 (2.5 μM) on day six for 24h. Whole cell lysates were subjected to SDS-PAGE and probed with the indicated antibodies. **F.** *Top panel:* Western blot analysis of EPAC2 overexpression (EPAC2-OE). *Bottom left panel:* Effect of EPAC2-OE on senescence in $BRAF^{V600E}$ transformed melanocytes. *Bottom right panel:* Effect of EPAC2-OE on the growth of transformed melanocytes.

**Supplementary Figure. 3: Systemic inhibition of *Epac* activity delayed melanomagenesis and tumor growth in *Braf/Pten (BPT)* mouse melanoma model.** Photographs of *BPT* mice five weeks after treatment with vehicle (control) and ESI-09 (EPACi).

**Supplementary Figure. 4: Genotyping of *BPT-Epac1/2* null mice tumor tissues. A.** Genomic PCR analysis of *BPT*, *BPT-Epac1/2* heterozygous and homozygous tumor tissues. The anticipated size of the respective wild type (WT) and knockout (KO) alleles of *Epac1* and *Epac2* are shown. **B** and **C.** Genotyping primer sequences and PCR conditions.

**Supplementary Figure. 5: Conditional deletion of *Epac1* or *Epac2* alone did not affect tumor growth and survival. A.** Representative photograph of *BPT*, $Epac1^{-/-}$ *and* $Epac2^{-/-}$ mice showing differences in tumor development at week five after 4HT application. **B.** The red circles represent tumors on the flanks of *BPT,* $Epac1^{-/-}$ *and* $Epac2^{-/-}$ mice at week five after 4HT application. **C.** Melanoma tumor latency (days) in *BPT,* $Epac1^{-/-}$ *and* $Epac2^{-/-}$ mice. Data are shown as mean ± SD, analyzed using One-way ANOVA using Tukey multiple comparison test and p-values are shown. **D.** Comparison of tumor incidence between *BPT,* $Epac1^{-/-}$ *and* $Epac2^{-/-}$ mice. n = number of tumors induced in each group. Two tumors were induced per mouse (each flank one tumor). Tumor incidence is presented as percent incidence. Data are shown as mean ± SD, analyzed using One-way ANOVA using Tukey multiple comparison test and p-values are shown. **E.** Tumor burden in *BPT,* $Epac1^{-/-}$ *and* $Epac2^{-/-}$ mice. Each line in the plot represents tumor burden in individual mouse. **F.** Kaplan-Meier survival analysis of *BPT, Epac1*$^{-/-}$ *and Epac2*$^{-/-}$ mice. Log-rank (Mantel-Cox) test was performed for survival trends between experimental groups. **G.** Table showing median survival and hazard ratio values between experimental groups. The median survival, hazard ratio, and CI were determined from the Kaplan-Meier survival curve and log-rank (Mantel-Cox) test.

**Supplementary Figure. 6: Deletion of *Epac1/2* reduced tumor growth and improved overall survival in mice. A.** Photographs of *BPT, Epac1/2*-double heterozygous and double homozygous mice showing differences in tumor development at week five after 4HT application. **B.** Hematoxylin and eosin (H&E) and S100A staining in *BPT*, *Epac1/2*-heterozygous and homozygous mice tumor sections.

**Supplementary Figure. 7: EPAC and the role of immune response in melanoma growth. A.** Fluorescence microscope images of *BPT, Epac1/2*-double heterozygous and double homozygous mice tumor sections stained for CD8 and CD4 T cell markers. Scale bar: 100 µm. **B**. Percentage of CD8 positive cells in *BPT, Epac1/2*-double heterozygous and double homozygous mice tumor sections. Data are shown as mean ± SD and analyzed using One-way ANOVA using Tukey multiple comparison test and p-values are shown. **C.** Schematic for treatment regimen of immunocompromised NSG mice xenografted with patient primary (WM115) and metastatic (MRA6) melanoma cells (BioRender). **D** and **E.** Photographs of tumors three weeks after treatment of NSG mice bearing WM115 and MRA6 tumors with indicated drugs.

**Supplementary Figure. 8: Gene lists for the top five enriched pathways (A) and PCR and western blot validation of TXNIP and ARRDC4. A.** List of most significantly up and downregulated genes from top five enriched pathways in MSIgDB hallmark database. **B.** qRT-PCR analysis of basal TXNIP mRNA expression in a panel of primary and metastatic melanoma cell lines. **C.** qRT-PCR validation of downregulation of TXNIP in a panel of primary and metastatic melanoma cell lines treated with ESI-09 for 12h. GAPDH is used as an internal control. Fold change in TXNIP expression determined by using $2^{-\Delta\Delta Ct}$ method. **D.** qRT-PCR analysis of basal ARRDC4 mRNA expression in a panel of primary and metastatic melanoma cell lines. **E.** qRT-PCR validation of downregulation of ARRDC4 in a panel of primary and metastatic melanoma cell lines treated with ESI-09 for 12h. GAPDH is used as an internal control. Fold change in ARRDC4 expression determined by using $2^{-\Delta\Delta Ct}$ method. **F.** Western blot analysis of the effect of ESI-09 treatment (24h) on ARRDC4 expression using a panel of primary and metastatic melanoma cell lines (**D:** DMSO; **E:** ESI-09).

**Supplementary Figure. 9: Genotyping of *Epac1*$^{+/-}$*/Epac2*$^{+/-}$ and *Epac1*$^{-/-}$ */Epac2*$^{-/-}$ tumor cell lines:** **A**. Genomic PCR analysis of tumor cell lines derived from *BPT*, *Epac1/2* heterozygous and homozygous mice tumor tissues. The anticipated size of the respective wild type (WT) and knockout (KO) alleles of *Epac1* and *Epac2* are shown. **B.** Western blot validation of oncogenic *Braf*$^{V600E}$expression and *Pten* loss in tumor cell lines. GAPDH is used as loading control.

**Supplementary Figure. 10: Effect of ESI-09 on various transcription factors known to regulate TXNIP expression. A.** Fold change in mRNA transcripts for various transcription factors from RNAseq analysis. **B.** Western blot analysis of MondoA, FoxO1, FoxO3, NRF2 in primary and metastatic melanoma cell lines treated with ESI-09 for 24h. GAPDH or β-tubulin is used as a loading control. **C.** qRT-PCR validation of upregulated HIF-1A expression using a panel of primary and LN-met melanoma cells treated with DMSO or ESI-09 for 12h. GAPDH is used as internal control. Fold change in HIF-1A expression determined by using $2^{-\Delta\Delta Ct}$ method. Data are presented as mean ± SD, analyzed using Unpaired t-test and p-values are shown. **D. .** Western blot validation of HIF-1α expression using a panel of primary and metastatic melanoma cell lines treated with DMSO or ESI-09 for 72h (**D:** DMSO; **E:** ESI-=09). **E.** Western blot analysis of HIF-1α and TXNIP expression changes in WM115 primary melanoma cells treated with ESI-09 or $CoCl_2$ alone or both for 12h.

**Supplementary Figure. 11: Effect of mitochondrial prooxidant rotenone on the growth of human melanoma cells. A.** Primary (WM1862, WM115), LN-met (WM165-1), and distant met (MRA5) cells were plated in six replicates in a 96-well plate (5000 cells/well) and treated with ESI-09 (2.5 μM) or rotenone (1 μM) alone or both for 72h. Cell numbers were estimated using MTT assay. Data are presented as mean ± SD, analyzed using One-way ANOVA with Tukey multiple comparison test and p-values are shown. **B.** Intracellular ROS levels were assessed using CellROX Green staining in above mentioned primary and metastatic melanoma cells treated with ESI-09 or GSH for 24h. Green fluorescence was visualized using fluorescence microscope. Scale bar: 200 μm. **C**. The graph shows changes in fluorescence intensity across different treatment conditions in primary and metastatic melanoma cell lines. Fluorescence intensity quantified using ImageJ V1.54S (NIH, US). Data are shown as mean ± SD and analyzed using One-way ANOVA using the Tukey multiple comparison test, and p-values are shown.

**Supplementary Figure. 12: EPAC signaling does not promote phenotype switch of primary melanoma. A.** Western blot analysis of changes in phenotype switch markers in patient matched primary, LN-met and unmatched primary and metastatic melanoma cells treated with DMSO or ESI-09 (2.5 μm) for 24h. **B** and **C.** Cell migration was analyzed by wound healing assay in a panel of primary and metastatic melanoma cells treated with DMSO or ESI-09 (2.5 μM) at 0, 6, 24, and 48 h. **D-F.** Cell invasion was analyzed by Matrigel invasion chambers in a panel of primary and metastatic melanoma cells treated with DMSO or ESI-09 (2.5 μM) for 48h.

### Figure. S1

A

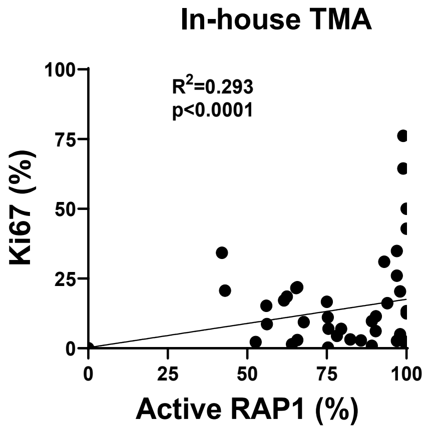


B

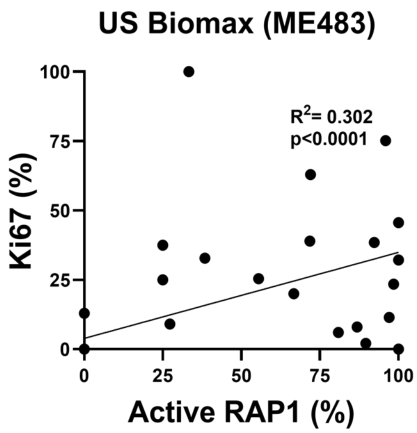


C

**Clinicopathological characteristics of in-house TMA**

| Primary Melanoma | N = 43 |
|---|---|
| **Clinical parameters** | **N (%)** |
| **Age** | |
| < 40 yrs | 13 (30) |
| 41-50 yrs | 5 (12) |
| 51-60 yrs | 6 (14) |
| 61-70 yrs | 13 (30) |
| 71-90 yrs | 6 (14) |
| **Tumour staging** | |
| Tis | 5 (12) |
| T1 | 34 (79) |
| T2 | 2 (5) |
| T3 | 1 (2) |
| T4 | 1 (2) |
| **Sex** | |
| Male | 29 (67) |
| Female | 14 (33) |
| **Breslow Thickness** | |
| ≤ 1mm | 39 (91) |
| > 1-2mm | 2 (5) |
| >2-4 mm | 2 (5) |

D

**Univariate Cox Regression Analyses of Disease-Free Survival**

| Clinicopathological parameters | Hazard ratio (95% CI) | p-value |
|---|---|---|
| Age (>40yrs vs <40yrs) | 0.7 (0.15-3.1) | 0.51 |
| Sex (Male vs Female) | 1.2 (0.26-5.41) | 0.82 |
| Breslow thickness (≥ 1-4mm vs <1mm) | 2.9 (0.34-25.25) | 0.33 |
| Tumour staging (T2-T4 vs Tis-T1) | **7.0 (1.3-38.34)** | **0.026** |
| RAP1-GTP expression (High vs Low) | **8.01 (0.91-70.28)** | **0.023** |

E

**Clinicopathological characteristics of Biomax TMA ME483**

| Primary Melanoma (32) | | Metastatic Melanoma (16) | |
|---|---|---|---|
| **Clinical parameters** | **N (%)** | **Clinical parameters** | **N (%)** |
| **Age** | | **Distant organ** | |
| < 40 yrs | 2 (6) | **Lymph Node** | 16 |
| 41-50 yrs | 12 (38) | **Age** | |
| 51-60 yrs | 9 (28) | < 40 yrs | 3 (19) |
| 61-70 yrs | 5 (16) | 41-50 yrs | 7 (44) |
| 71-90 yrs | 4 (13) | 51-60 yrs | 2 (13) |
| **Tumour staging** | | 61-70 yrs | 3 (19) |
| Tis | - | 71-90 yrs | 1 (6) |
| T1 | - | **Sex** | |
| T2 | 29 (91) | Male | 8 (50) |
| T3 | 3 (9) | Female | 8 (50) |
| T4 | - | | |
| **Sex** | | | |
| Male | 21 (66) | | |
| Female | 11 (34) | | |

### Figure. S2

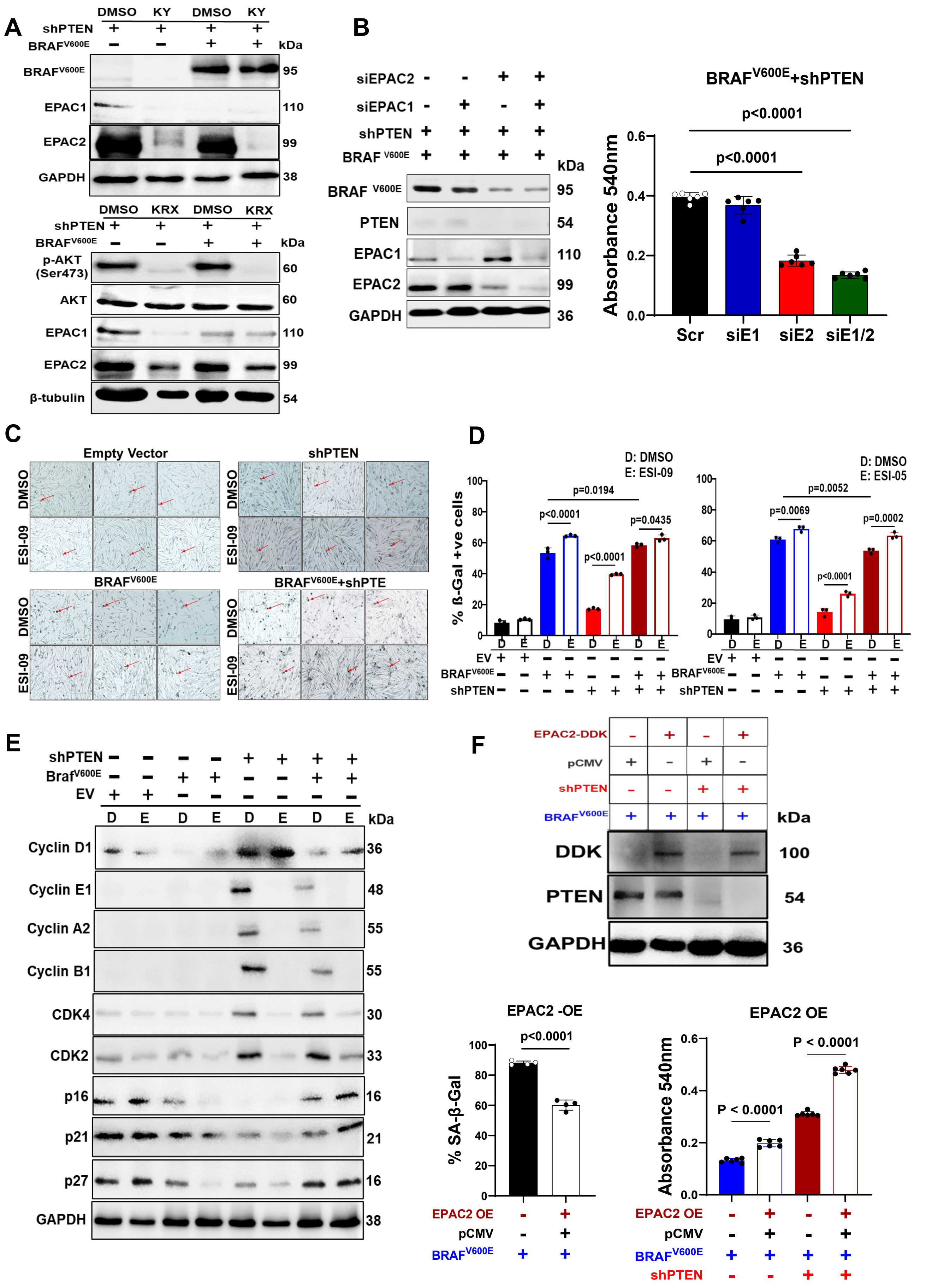

A
DMSO KY DMSO KY
shPTEN
BRAFV600E
kDa
BRAFV600E 95
EPAC1 110
EPAC2 99
GAPDH 38
DMSO KRX DMSO KRX
p-AKT (Ser473) 60
AKT 60
EPAC1 110
EPAC2 99
β-tubulin 54
B
siEPAC2
siEPAC1
shPTEN
BRAF V600E
PTEN 54
EPAC2 99
GAPDH 36
BRAFV600E+shPTEN
p<0.0001
Absorbance 540nm
Scr siE1 siE2 siE1/2
C
Empty Vector
shPTEN
DMSO
ESI-09
BRAFV600E
BRAFV600E+shPTE
D
D: DMSO
E: ESI-09
E: ESI-05
% ß-Gal +ve cells
p=0.0194
p=0.0435
p=0.0052
p=0.0069
p=0.0002
EV
E
Braf V600E
Cyclin D1 36
Cyclin E1 48
Cyclin A2 55
Cyclin B1 55
CDK4 30
CDK2 33
p16 16
p21 21
p27 16
F
EPAC2-DDK
pCMV
DDK 100
EPAC2 -OE
% SA-β-Gal
EPAC2 OE
P < 0.0001

### Figure. S3

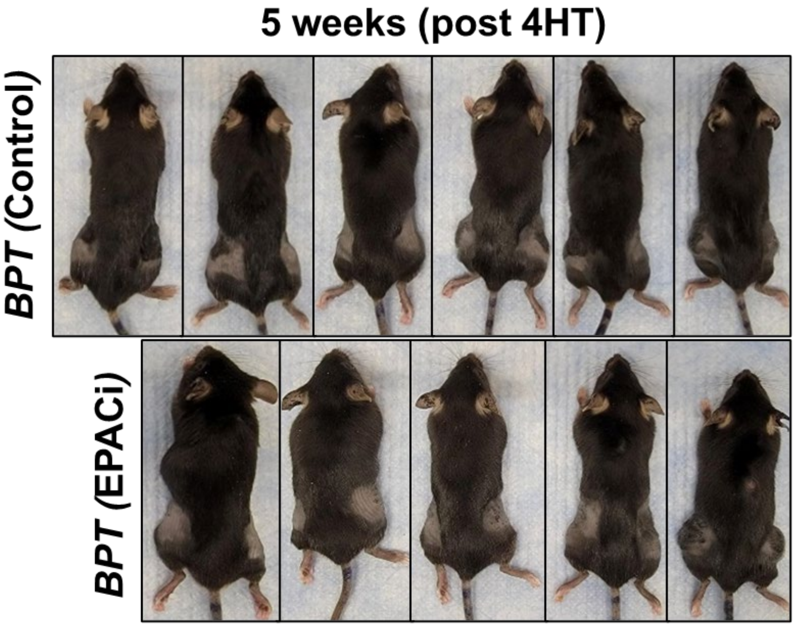

### Figure. S4

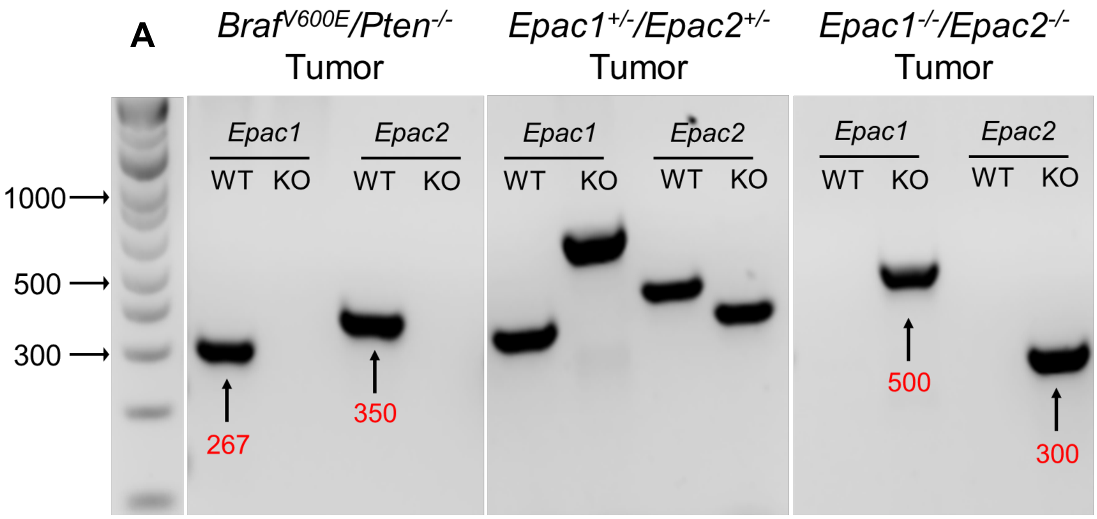


**B**

Genotyping primer sequences:

| | |
|---|---|
| Epac1 WT | 5'-CTGGCCTCTCCTGAATCTTG-3'<br>5'-CCTCGCTGTTGGTAAGTGGT-3' |
| Epac1 KO | 5'-AATGGGCTGACCGCTTCCTCGT-3'<br>5'-GCCATAGCCTCAACAAGCTC-3' |
| Epac2 WT | 5'-CCTCCCTTTTGCTCTCTCCT-3'<br>5'-CGCTCGCTGCATTTGTATTA-3' |
| Epac2 KO | 5'-AATGGGCTGACCGCTTCCTCGT-3'<br>5'-CGCTCGCTGCATTTGTATTA-3' |

**C**

PCR conditions:

94ºC/5 min
94ºC/30 sec
60ºC/30 sec | 30 cycles
72ºC/30 sec
72ºC/10 min
4ºc/∞

### Figure. S5

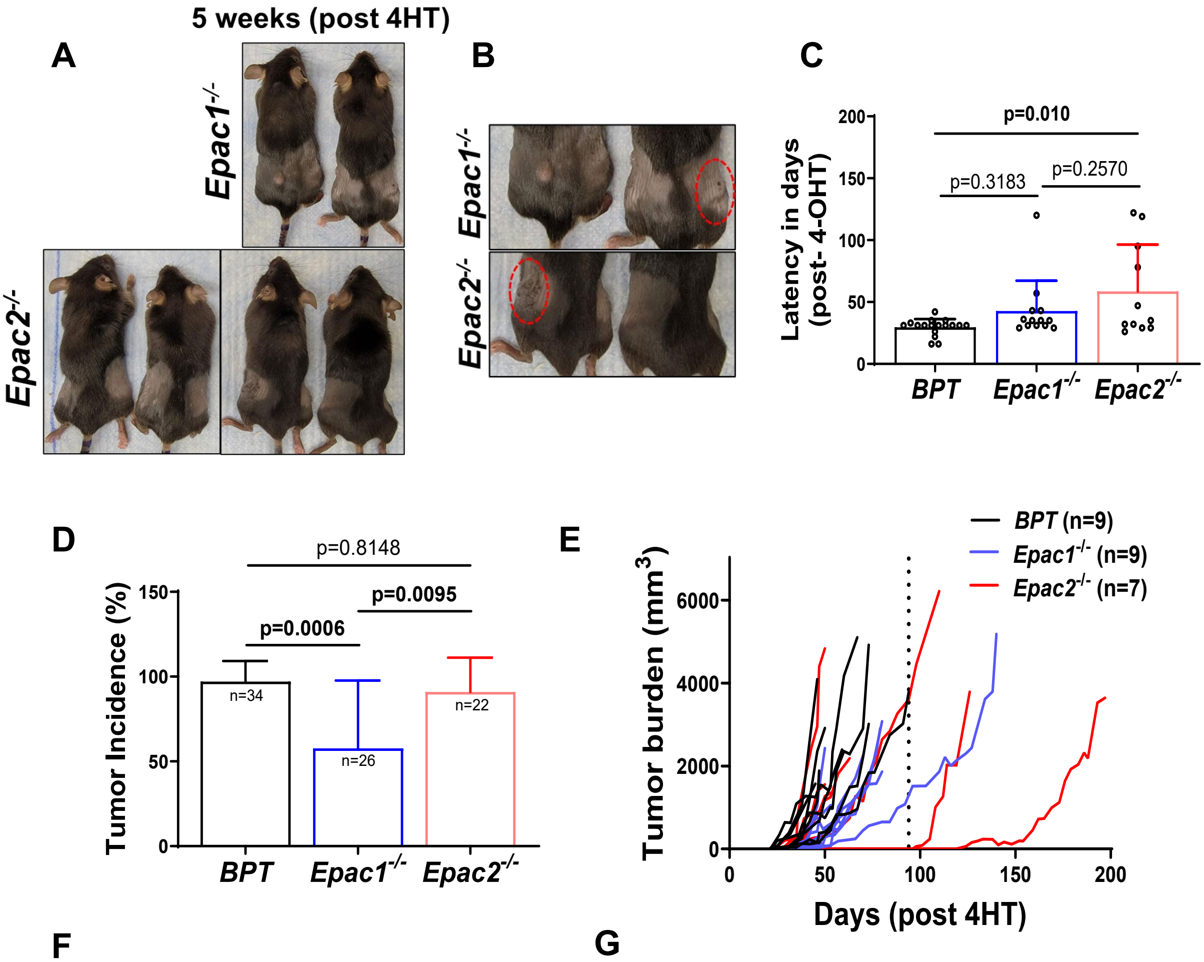


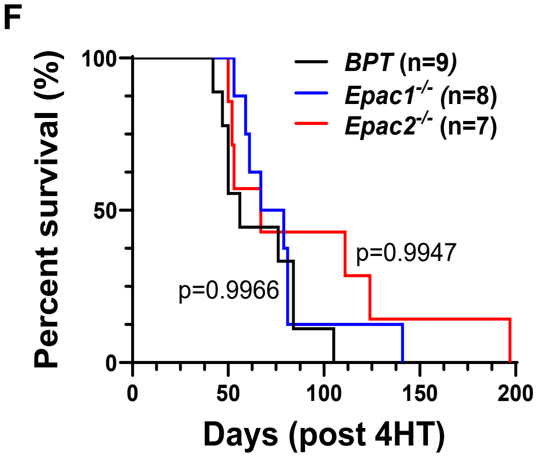


G

| Genotype | Median Survival (days) | Hazard Ratio |
|---|---|---|
| *BPT* | 56 | - |
| *Epac1-/-* | 73 | 0.7718 (95% CI: 0.297 – 2.002) |
| *Epac2-/-* | 67 | 0.5222 (95% CI: 0.192 – 1.419) |

### Figure. S6

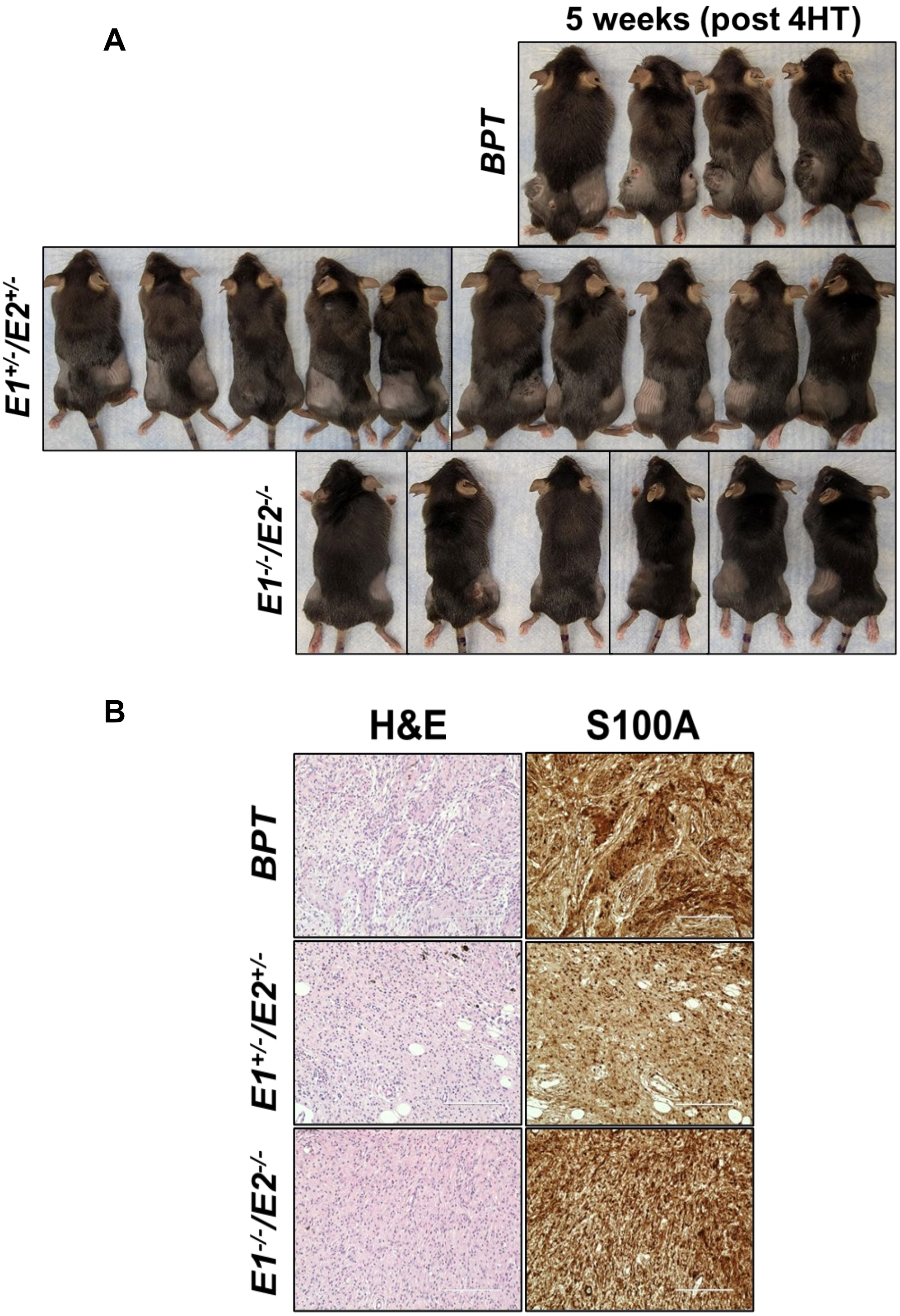

### Figure. S7

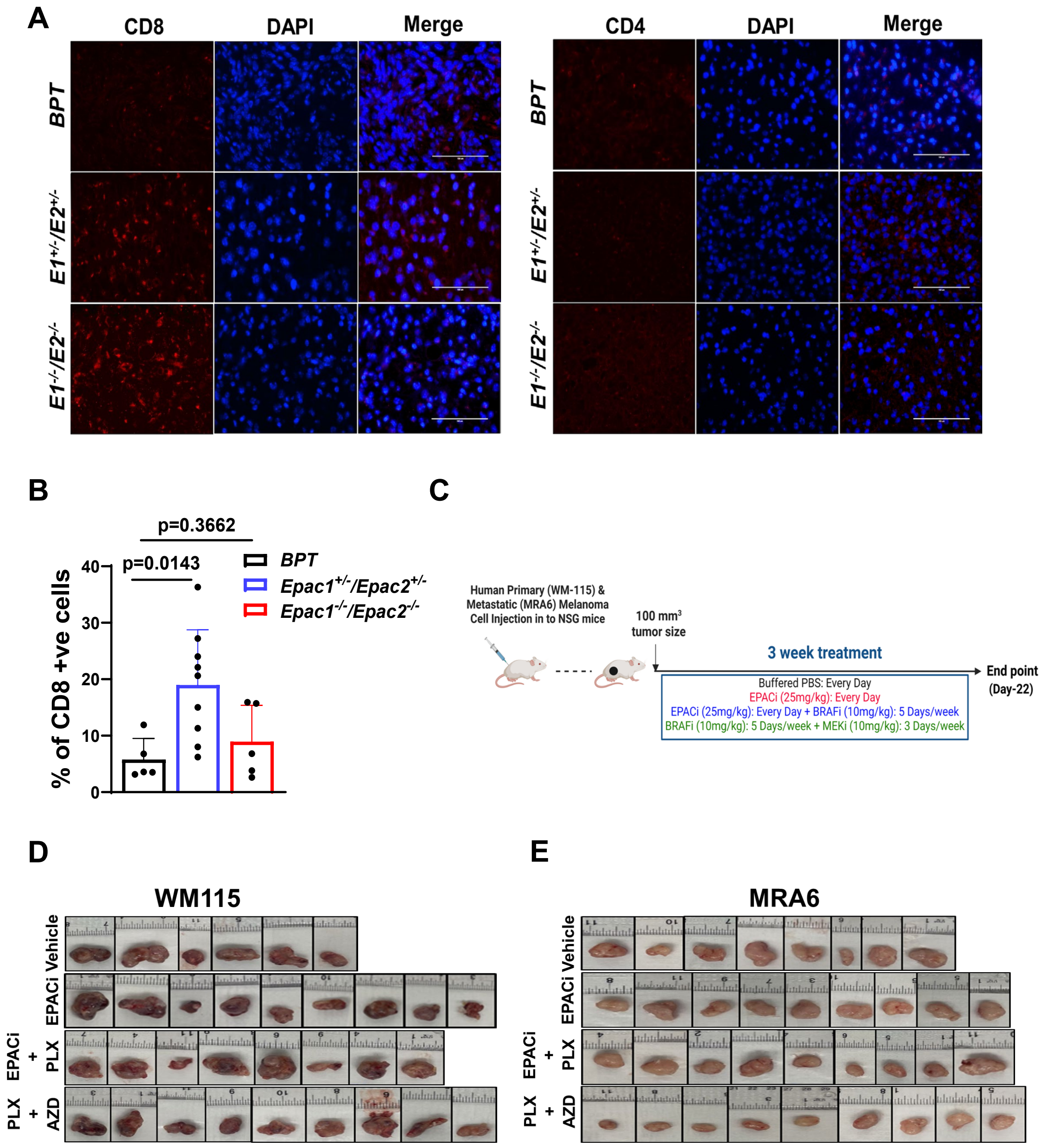
A
CD8
DAPI
Merge
CD4
DAPI
Merge
BPT
E1+/-/E2+/-
E1-/-/E2-/-
B
p=0.3662
p=0.0143
% of CD8 +ve cells
0
10
20
30
40
BPT
Epac1+/-/Epac2+/-
Epac1-/-/Epac2-/-
C
Human Primary (WM-115) & Metastatic (MRA6) Melanoma Cell Injection in to NSG mice
100 mm3 tumor size
3 week treatment
End point (Day-22)
Buffered PBS: Every Day
EPACi (25mg/kg): Every Day
EPACi (25mg/kg): Every Day + BRAFi (10mg/kg): 5 Days/week
BRAFi (10mg/kg): 5 Days/week + MEKi (10mg/kg): 3 Days/week
D
WM115
Vehicle
EPACi
EPACi + PLX
PLX + AZD
E
MRA6
Vehicle
EPACi
EPACi + PLX
PLX + AZD

### Figure. S8

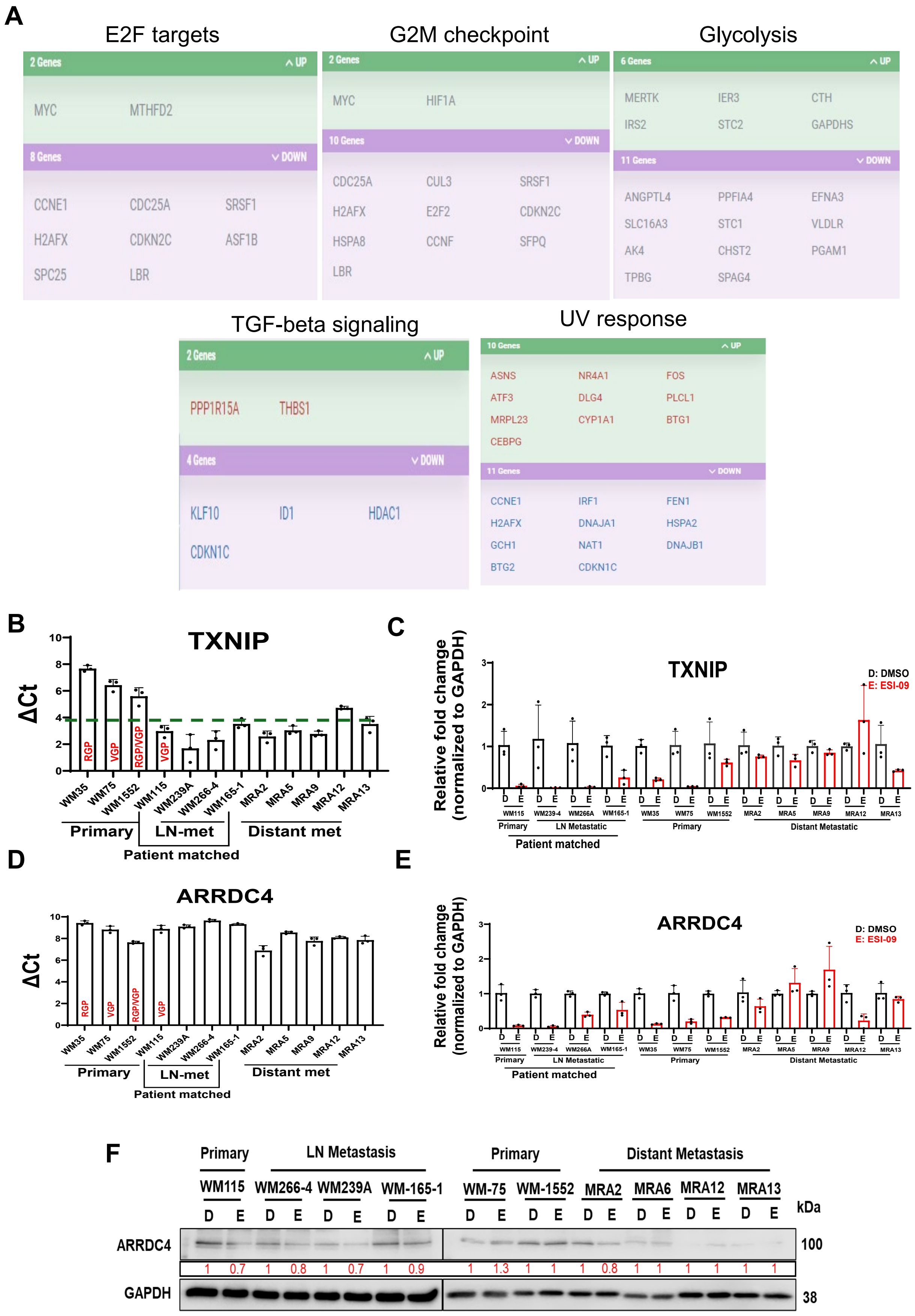
A
E2F targets
2 Genes UP
MYC MTHFD2
8 Genes DOWN
CCNE1 CDC25A SRSF1
H2AFX CDKN2C ASF1B
SPC25 LBR
G2M checkpoint
2 Genes UP
MYC HIF1A
10 Genes DOWN
CDC25A CUL3 SRSF1
H2AFX E2F2 CDKN2C
HSPA8 CCNF SFPQ
LBR
Glycolysis
6 Genes UP
MERTK IER3 CTH
IRS2 STC2 GAPDHS
11 Genes DOWN
ANGPTL4 PPFIA4 EFNA3
SLC16A3 STC1 VLDLR
AK4 CHST2 PGAM1
TPBG SPAG4
TGF-beta signaling
2 Genes UP
PPP1R15A THBS1
4 Genes DOWN
KLF10 ID1 HDAC1
CDKN1C
UV response
10 Genes UP
ASNS NR4A1 FOS
ATF3 DLG4 PLCL1
MRPL23 CYP1A1 BTG1
CEBPG
11 Genes DOWN
CCNE1 IRF1 FEN1
H2AFX DNAJA1 HSPA2
GCH1 NAT1 DNAJB1
BTG2 CDKN1C
B
TXNIP
ΔCt
RGP VGP RGP/VGP VGP
WM35 WM75 WM1552 WM115 WM239A WM266-4 WM165-1 MRA2 MRA5 MRA9 MRA12 MRA13
Primary LN-met Distant met
Patient matched
C
TXNIP
Relative fold chamge (normalized to GAPDH)
D: DMSO
E: ESI-09
WM115 WM239-4 WM266A WM165-1 WM35 WM75 WM1552 MRA2 MRA5 MRA9 MRA12 MRA13
Primary LN Metastatic Primary Distant Metastatic
Patient matched
D
ARRDC4
ΔCt
RGP VGP RGP/VGP VGP
WM35 WM75 WM1552 WM115 WM239A WM266-4 WM165-1 MRA2 MRA5 MRA9 MRA12 MRA13
Primary LN-met Distant met
Patient matched
E
ARRDC4
Relative fold chamge (normalized to GAPDH)
D: DMSO
E: ESI-09
WM115 WM239-4 WM266A WM165-1 WM35 WM75 WM1552 MRA2 MRA5 MRA9 MRA12 MRA13
Primary LN Metastatic Primary Distant Metastatic
Patient matched
F
Primary LN Metastasis Primary Distant Metastasis
WM115 WM266-4 WM239A WM-165-1 WM-75 WM-1552 MRA2 MRA6 MRA12 MRA13
D E D E D E D E D E D E D E D E D E D E
kDa
ARRDC4 100
1 0.7 1 0.8 1 0.7 1 0.9 1 1.3 1 1 1 0.8 1 1 1 1 1 1
GAPDH 38

**Figure. S9**

**A**

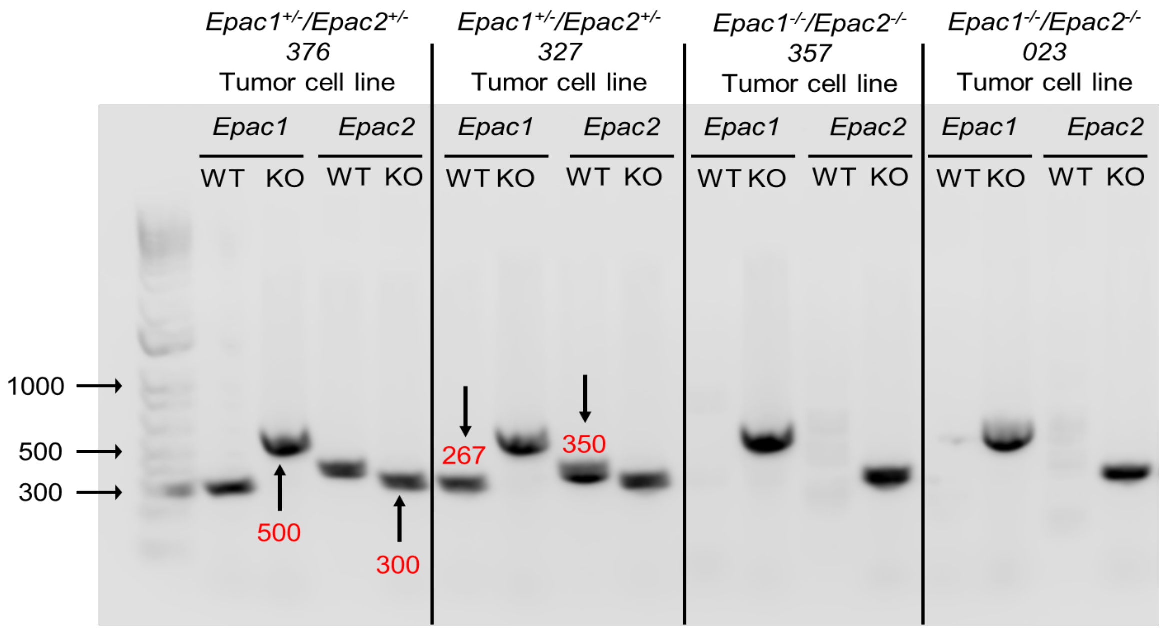

Epac1+/-/Epac2+/-
376
Tumor cell line
Epac1+/-/Epac2+/-
327
Tumor cell line
Epac1-/-/Epac2-/-
357
Tumor cell line
Epac1-/-/Epac2-/-
023
Tumor cell line
Epac1
Epac2
WT
KO
1000
500
300
267
350


**B**

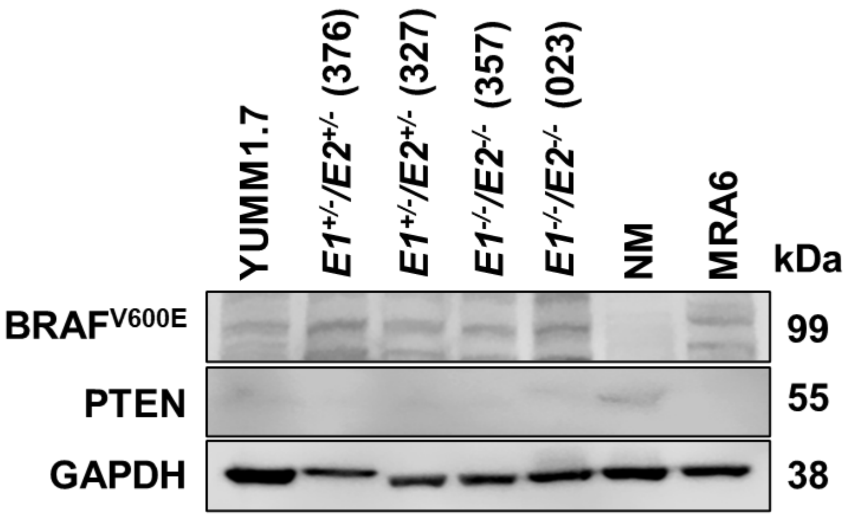

YUMM1.7
E1+/-/E2+/- (376)
E1+/-/E2+/- (327)
E1-/-/E2-/- (357)
E1-/-/E2-/- (023)
NM
MRA6
kDa
BRAFV600E
PTEN
GAPDH
99
55
38

### Figure. S10

A

| Log2 fold change with p value | | | |
|---|---|---|---|
| Gene | Treatment | WM115 | WM165 |
| MONDOA | 6h | -0.013 (p=0.829) | -0.001 (p=0.979) |
| | 12h | 0.018(p=0.824) | 0.109 (p=0.318) |
| FoxO1 | 6h | -0.059(p=0.438) | 0.092 (p=0.165) |
| | 12h | 0.834 (p=8.46E-16) | -0.106 (p=0.354) |
| FoxO3 | 6h | -0.090(p=0.135) | -0.017 (p=0.745) |
| | 12h | 0.355 (p=2.43E-07) | -0.008 (p=0.932) |
| HIF-1A | 6h | -0.088 (p=0.126) | -7.10E-05 (p=0.998) |
| | **12h** | **1.215 (p=1.13E-59)** | 0.015 (p=0.854) |
| NFE2L2 | 6h | -0.040 (p=0.490) | -0.165 (p=0.670) |
| | 12h | 0.662 (p=1.76E-21) | -0.096 (p=0.200) |

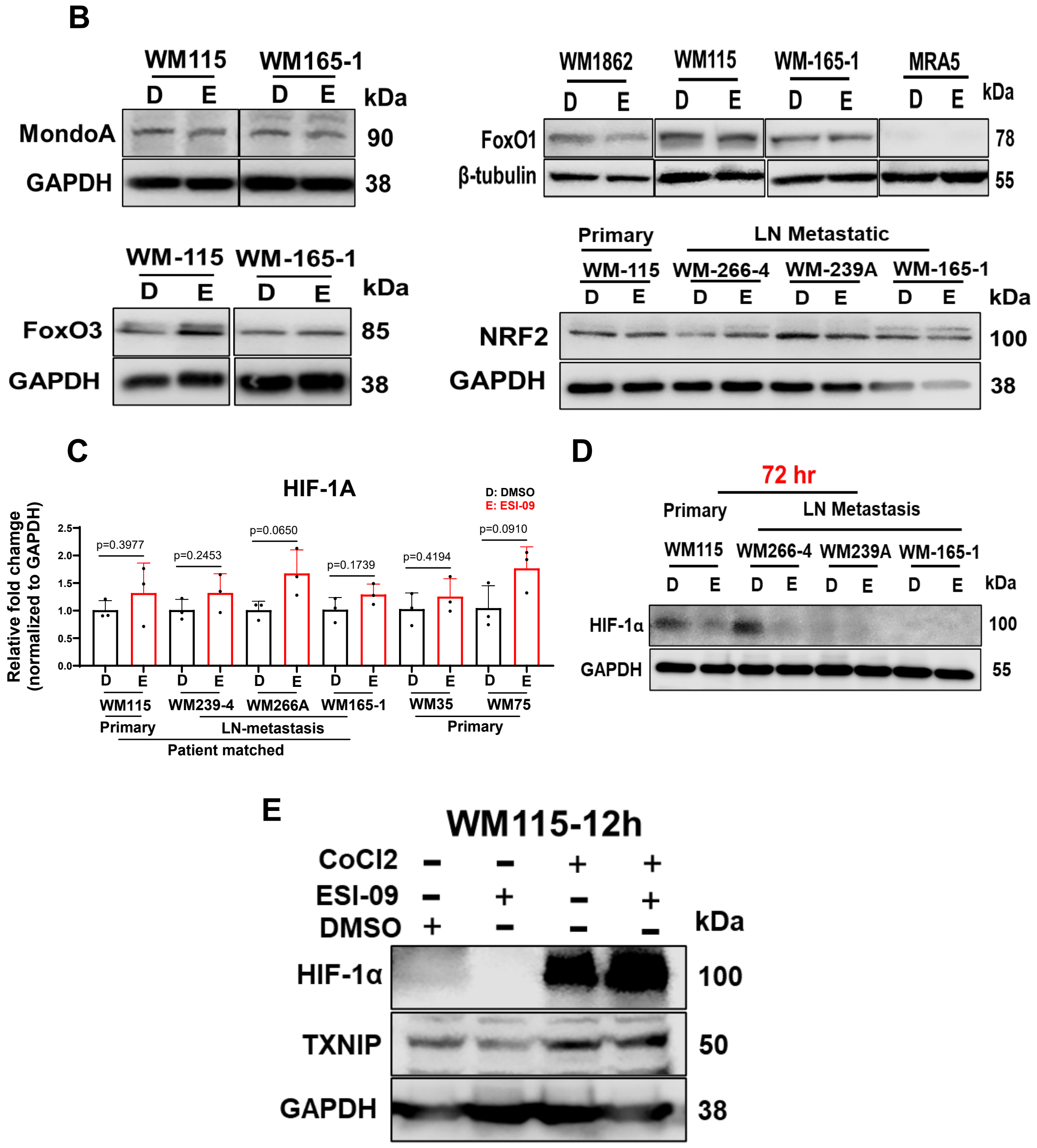

### Supplementary Figure. 11

A

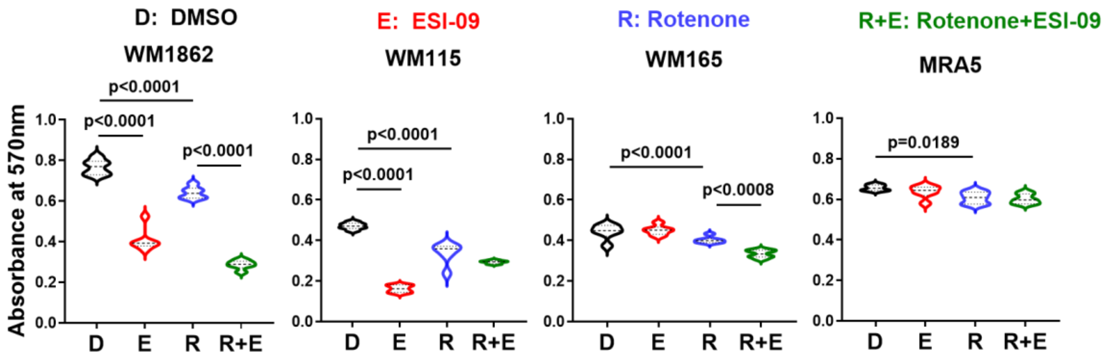


B

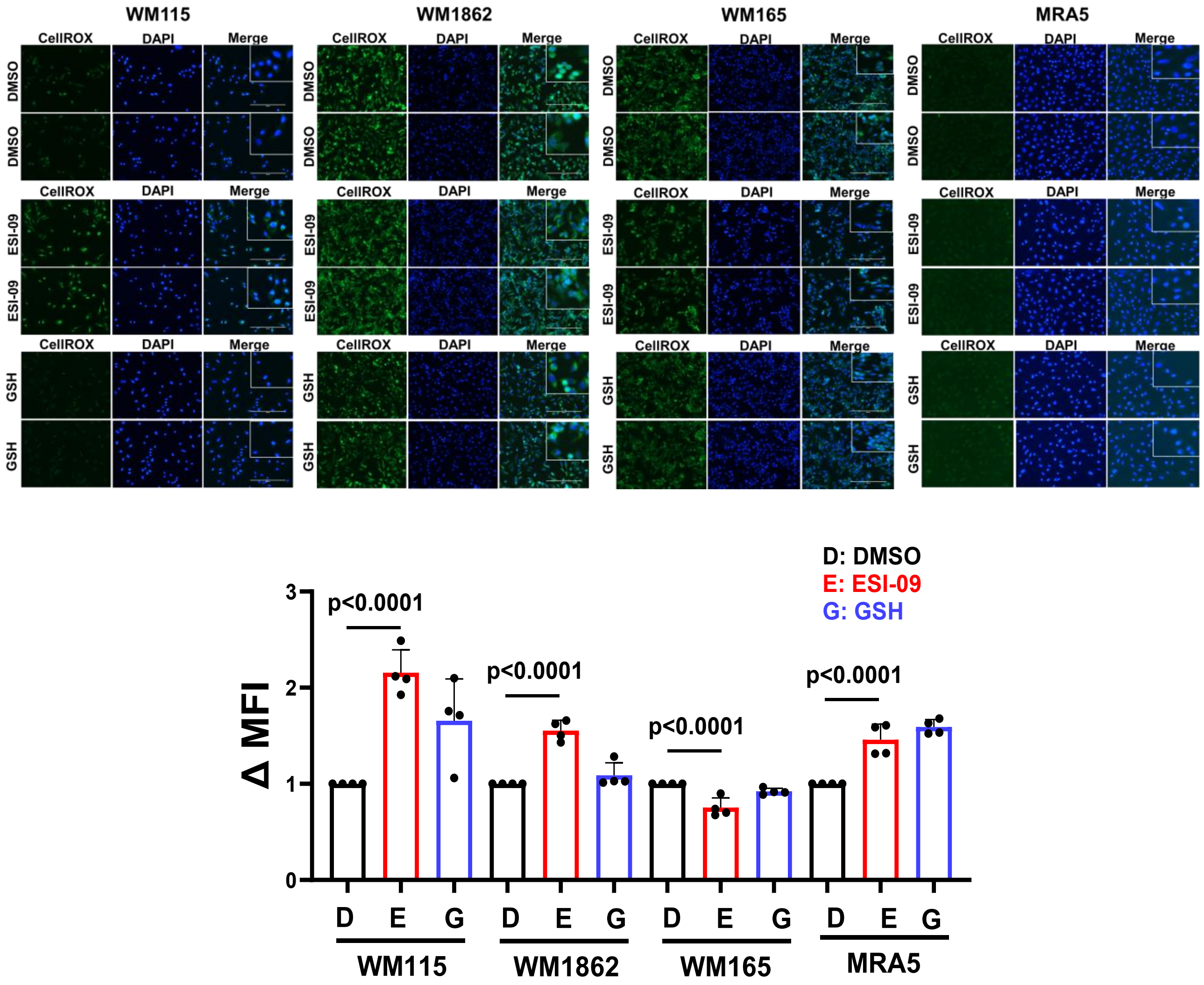

### Figure. S12

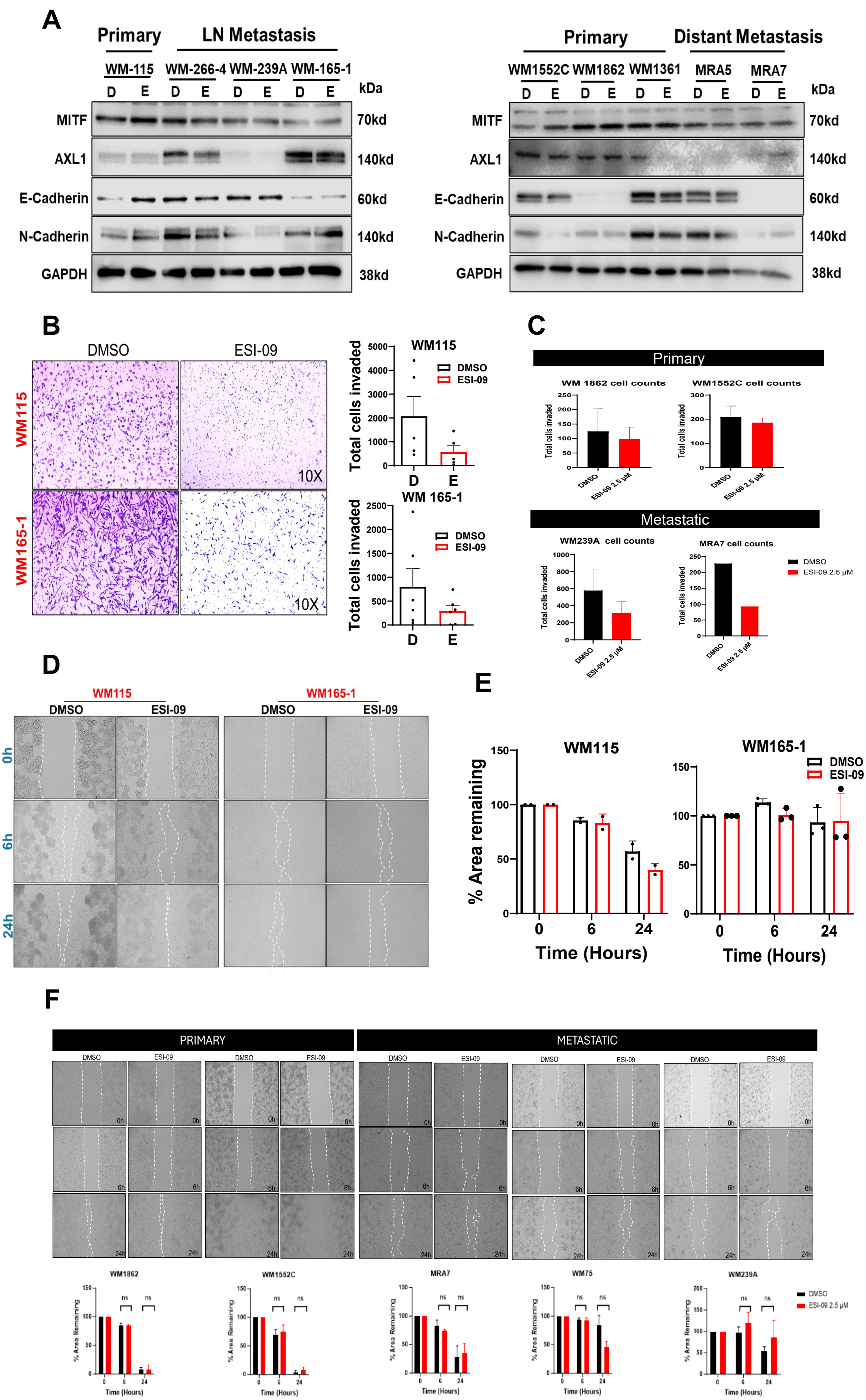
A
Primary
LN Metastasis
WM-115
WM-266-4
WM-239A
WM-165-1
D E D E D E D E
kDa
MITF
AXL1
E-Cadherin
N-Cadherin
GAPDH
70kd
140kd
60kd
140kd
38kd
Primary
Distant Metastasis
WM1552C
WM1862
WM1361
MRA5
MRA7
B
DMSO
ESI-09
WM115
WM165-1
10X
Total cells invaded
WM115
WM 165-1
DMSO
ESI-09
D
E
C
Primary
WM 1862 cell counts
WM1552C cell counts
Metastatic
WM239A cell counts
MRA7 cell counts
Total cells invaded
DMSO
ESI-09 2.5 μM
D
0h
6h
24h
E
% Area remaining
Time (Hours)
F
PRIMARY
METASTATIC
WM1862
WM1552C
MRA7
WM75
WM239A
% Area Remaining
ns
Time (Hours)
